# Action space restructures visual working memory in prefrontal cortex

**DOI:** 10.1101/2023.08.13.553135

**Authors:** Donatas Jonikaitis, Shude Zhu

## Abstract

Visual working memory enables flexible behavior by decoupling sensory stimuli from behavioral actions. While previous studies have predominantly focused on the storage component of working memory, the role of future actions in shaping working memory remains unknown. To answer this question, we used two working memory tasks that allowed the dissociation of sensory and action components of working memory. We measured behavioral performance and neuronal activity in the macaque prefrontal cortex area, frontal eye fields. We show that the action space reshapes working memory, as evidenced by distinct patterns of memory tuning and attentional orienting between the two tasks. Notably, neuronal activity during the working memory period predicted future behavior and exhibited mixed selectivity in relation to the sensory space but linear selectivity relative to the action space. This linear selectivity was achieved through the rapid transformation from sensory to action space and was subsequently maintained as a stable cross-temporal population activity pattern. Combined, we provide direct physiological evidence of the action-oriented nature of frontal eye field neurons during memory tasks and demonstrate that the anticipation of behavioral outcomes plays a significant role in transforming and maintaining the contents of visual working memory.

## Introduction

A key characteristic of primate behavior is its remarkable flexibility. This flexibility extends beyond reflexive behavior and, depending on context and goals, allows for different responses to identical sensory inputs. Visual working memory supports this flexibility, as it enables the storage and manipulation of sensory information that is no longer available (Baddeley, 1992; Funahashi, 2022; Gazzaley & Nobre, 2012; Ma et al., 2014; Myers et al., 2017). However, despite the crucial role of visual working memory in guiding future behavior, past neurophysiological studies have frequently focused solely on the storage of sensory information, as if it were independent from anticipated actions. Therefore, the question of whether and how the storage of sensory information interacts with anticipated actions remains unresolved.

Activity of the prefrontal cortex has often been the focus of physiological studies investigating neuronal correlates of visual working memory (Cavanagh et al., 2018; Clark et al., 2012; Funahashi et al., 1989; Fuster, 1973; Murray et al., 2017; Stokes et al., 2013; Umeno & Goldberg, 2001). On one hand, behavior relies on the accurate storage of sensory information; thus, research has investigated visual working memory as storage resembling past sensory inputs or “sensory space,” such as locations or features (Bays & Husain, 2008; Harrison & Tong, 2009; Kerkoerle et al., 2017; Wilken & Ma, 2004). In this case, memory is assumed to store accurate sensory information, with neuronal activity in the prefrontal cortex correlated with this information (Constantinidis et al., 2001; Ester et al., 2015; Mendoza-Halliday et al., 2014; Panichello & Buschman, 2021; Rezayat et al., 2021; Wasmuht et al., 2018; Xie et al., 2022). The implicit assumption in this approach is the independence of working memory from future actions.

On the other hand, the fundamental function of working memory is to link sensory inputs to motor actions. It has been proposed that sensory inputs automatically activate a repertoire of potential actions (Engel et al., 2013; Hommel et al., 2001), and consequently, visual working memory should also be considered in this context (Cisek & Kalaska, 2004; Ede, Chekroud, & Nobre, 2019; Hanning et al., 2016; Henderson et al., 2022; Lawrence et al., 2001; Ohl & Rolfs, 2017). Indeed, activity in the prefrontal cortex is also modulated by anticipated actions (Funahashi et al., 1993; Kastner et al., 2007; Khanna et al., 2019; Messinger et al., 2009; Sajad et al., 2016). In different experimental situations, this anticipation can incorporate response preparation for single (Funahashi et al., 1989; Sajad et al., 2016) or multiple parallel actions (Baldauf & Deubel, 2008b; Hanning et al., 2022), selection of some actions and inhibition of others (Hasegawa et al., 2004; Jonikaitis et al., 2019), and implementation of different stimulus-response rules (Everling & Munoz, 2000; Everling & DeSouza, 2005). To encompass this variety of behavioral strategies, we refer to it here as the “action space” (Baldauf & Deubel, 2010).

However, separating the contributions of sensory and action space to working memory has been surprisingly difficult, especially in spatial working memory tasks. The main reason for this is that these contributions frequently overlap (Jonikaitis & Moore, 2019). In a typically used delayed-response task, this overlap is complete, such as when making a motor action towards a memorized stimulus (Funahashi et al., 1989; Hanning et al., 2018; Heuer et al., 2017; Markowitz et al., 2015; Sajad et al., 2016). In such cases, studies have shown that spatial working memory, covert spatial attention, and action preparation all converge on the memorized location (Heuer et al., 2020; Jonikaitis & Moore, 2019). As a result, behavior and neuronal activity during the memory period could reflect either sensory or action space.

Alternatively, a second set of tasks attempted to separate motor actions from sensory stimuli. In the anti-saccade task, a motor action is directed to a location opposite from the sensory stimulus (Funahashi et al., 1993; Hallett, 1978; Zhang & Barash, 2000), and studies have reported neuronal responses tuned to the sensory stimulus and the anti-saccade locations (Funahashi et al., 1993; Saber et al., 2015; Steinmetz & Moore, 2014; M. Zhang & Barash, 2000). However, the relationship of this sensory and action-related activity to behavioral strategies in anti-saccade tasks is not known. Indeed, behavioral studies in humans have suggested that anti-saccade tasks involve competition between two motor actions - one to the sensory stimulus and one to the anti-saccade location (Noorani & Carpenter, 2013; Salinas et al., 2019) or shifts of covert spatial attention to both locations (Klapetek et al., 2016; Mikula et al., 2018).

In this study, we measured the contributions of sensory and action space to visual working memory using a set of tasks, which are uniquely suited for this question (Hasegawa et al., 2004; Jonikaitis et al., 2019, 2023). In two memory tasks, subjects have to memorize a location, but the expected action after the memory period is completely different. In one task - Look - memory location predicts motor action, and thus sensory and action spaces overlap. In the other task - Avoid - memory location does not predict motor action, and thus sensory and action spaces are dissociated. First, we employed a set of behavioral measures and manipulations to determine the behavioral strategies used throughout the memory period in Look and Avoid tasks. Next, we conducted simultaneous recordings of neuronal populations to measure activity in the frontal eye fields (FEF), a prefrontal cortical area consistently associated with spatial working memory and eye movements. Our findings demonstrate that instead of sensory space, action space has a stronger influence on behavioral performance and neuronal activity in the prefrontal cortical area FEF.

## Results

We trained two rhesus macaques to perform different versions of a spatial working memory task: Look, Avoid (Hanning et al., 2016; Jonikaitis et al., 2019, 2023), and memory-guided saccade (MGS) (Funahashi et al., 1989; Noudoost et al., 2021) tasks. On some experimental sessions, monkeys also completed a Fixation control task that did not require memory (Figure S1B). In all memory tasks, monkeys maintained their gaze on a central fixation point, and a briefly shown peripheral visual cue (50 ms) indicated a location to be memorized (Figure 1A and Figure S1A). In the Look and Avoid tasks, after the memory period, two choice targets appeared - one at the cued location and one at a non-cued, randomly selected location (Hanning et al., 2016; Hasegawa et al., 2004; Jonikaitis et al., 2019, 2023). In the Look task, monkeys were rewarded for making a saccade to the choice target at the cued and memorized location, whereas in the Avoid task - to the one at the non-cued and non-memorized location. This task design introduced a crucial dissociation between sensory and action space. Sensory space was matched in both tasks: monkeys had to memorize an identical set of locations, and accurate memory was necessary for correct task performance. Action space, however, was mismatched: while the cue predicted the saccade in the Look task (Jonikaitis et al., 2019; Lawrence et al., 2001), the cue only predicted a location to avoid, but not the saccade in the Avoid task (Jonikaitis et al., 2019).

**Figure 1.**
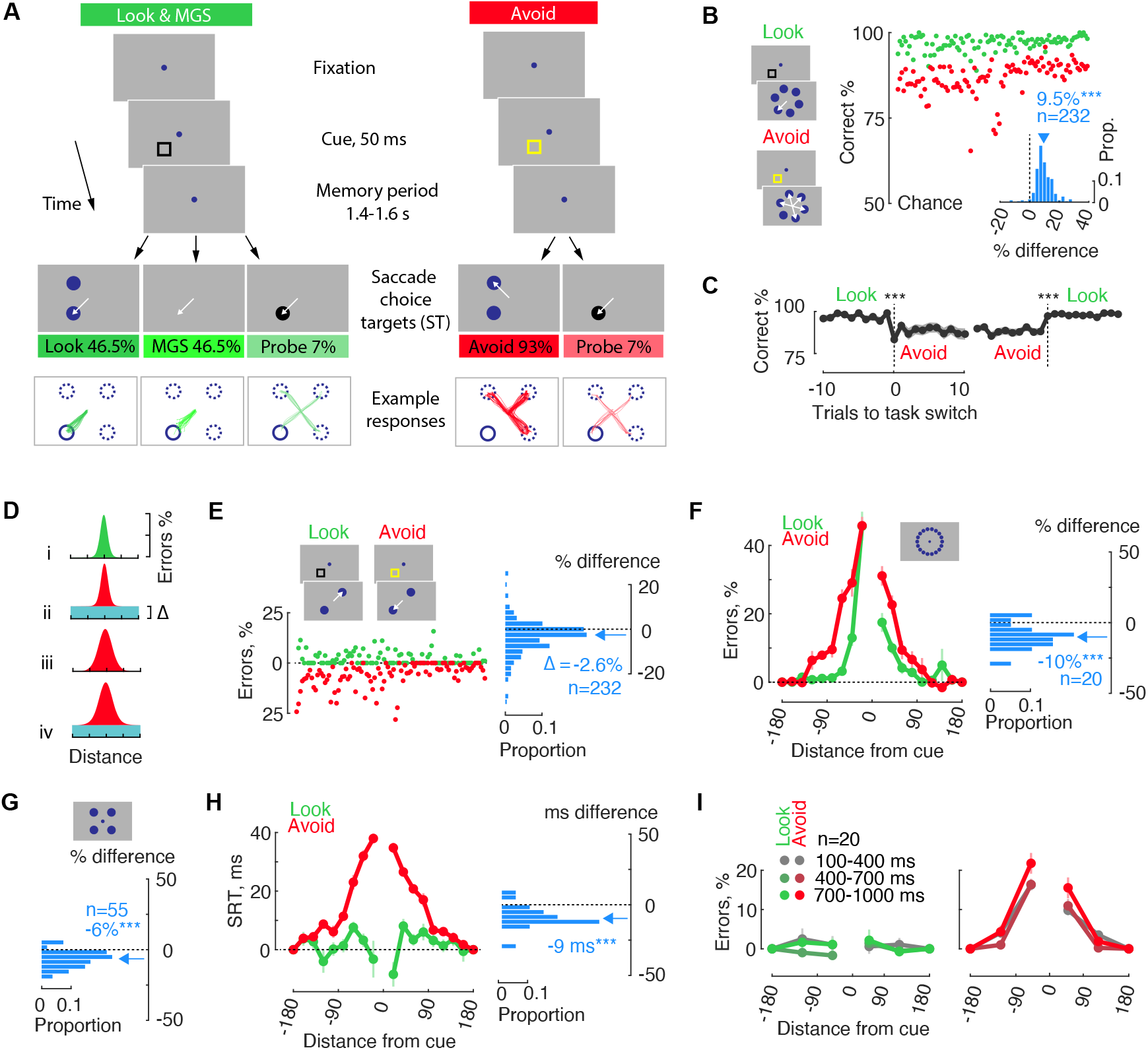
Task and performance. **(A)** Memory tasks. In all tasks, following central fixation, a briefly presented cue indicated the location to be memorized. In the Look task, after the delay, two saccade choice targets appeared: one at the cued location and another at a randomly selected location. Monkeys executed a saccadic eye movement to the target corresponding to the memorized cue location. In the MGS task, no targets were displayed, and monkeys directed their eye movement to the memorized location of the cue. Conversely, in the Avoid task, monkeys made an eye movement to a target located at a non-memorized, non-cued location. The cue color differed across tasks to emphasize the current rule. On a small proportion of trials—referred to as probe trials—a single target was presented, and monkeys executed an eye movement to the probe location. The bottom panels depict example eye movement trajectories from a FEF recording session. The trajectories are rotated in a way that the cue always appears at the lower left. **(B)** Memory performance in Look and Avoid tasks for a randomly selected subset of 100 sessions. The chance level is 50%. The inset shows a distribution of Look-Avoid performance differences for individual sessions. An arrow indicates the mean value across all sessions. Positive values indicate higher performance in the Look task. “n” denotes the number of sessions used. “**” indicates p<0.001; “” indicates p<0.05; “(ns)” signifies not significant. **(C)** Memory performance across task switches. Task switches occurred from Look to Avoid or vice versa at “Trial 0.” The figure displays 10 trials prior to and after the switch. The shaded areas indicate the standard error of the mean. A failure to switch the task would be indicated by a brief performance drop to 0%. **(D)** An illustration of the primary factors contributing to performance errors in Look and Avoid tasks. (i) Performance in the Look task can be understood as dependent on memory accuracy, represented by a spatial tuning function. This function is constructed from error trials based on the distance between response targets. (ii) A performance decrease in the Avoid task might arise from failure to adhere to the task rule, as evidenced by a uniform increase in errors across all saccade locations relative to the cue. This phenomenon preserves the tuning intact. (iii) Performance deterioration could stem from reduced memory accuracy, manifested as an increase in the width of the tuning function. (iv) A performance decrease might result from a combination of tuning changes and task rule failures. **(E)** A measure of failure to implement task rules. Performance errors are shown for trials where response targets were opposite each other. Histogram shows error probability difference between the Look and Avoid tasks. Negative values indicate more errors or more frequent failures to implement the appropriate task rule in the Avoid task. **(F)** An experiment involving an extended range of response target distances (inset displays possible cue locations, n=18) to evaluate memory tuning differences. Error rates are normalized relative to the location opposite from the cue to account for task rule failures. Histogram insets: % memory errors difference between Look and Avoid tasks for individual behavioral sessions. Negative values indicate more errors or a wider tuning function in the Avoid task. **(G)** The distribution of performance error probability differences during FEF recording sessions. **(H)** A tuning function constructed from saccadic reaction times on correct trials. **(I)** An experiment involving an extended range of delay durations, randomly selected from an interval of 100 to 1000 ms.

### Action space reshapes working memory tuning

First, we measured behavioral strategies in Look and Avoid tasks, which have been surprisingly difficult to measure in earlier studies. To accomplish this, we collected a comprehensive behavioral dataset (AQ: n=221 sessions; HB: n=190). Memory performance was higher in the Look task than in the Avoid task (monkey AQ: Look 97%, Avoid 87%, p<0.001; monkey HB: Look 81% and Avoid 72%, p<0.001) (Figure 1B and S3A). This performance difference aligns with studies that used a multi-task design, such as pro-saccades and anti-saccades in the same experimental session (Amador et al., 1998; Lebedev et al., 2004; Takeda & Funahashi, 2002). However, the reasons for this performance difference have not been addressed previously. First, we ruled out insufficient learning. Monkey AQ completed over 800 sessions (including training sessions), and monkey HB completed over 600 sessions throughout the multi-year experiment, during which the performance differences between the tasks were maintained (Figure 1B and S3A). Furthermore, robust performance was achieved during task switches. During each session, monkeys completed interleaved blocks of Look and Avoid tasks, and task-specific performance was observed immediately after the switch (Figure 1C, S2A, and S3B-C) (all p<0.001). Additionally, monkeys adjusted their performance after an error trial comparably frequently in Look and Avoid tasks (AQ: Look-Avoid difference −1.4±1.4%, p>0.05, HB: −1.1±1.1%, p>0.05). Lastly, differences between tasks cannot be explained by a speed-accuracy trade-off, as saccadic reaction times in the Avoid task were slower than in the Look task (Figure S2B and S3D) (AQ: Look 157 ms, Avoid 164 ms, p<0.001, HB: Look 176 ms, Avoid 184 ms, p<0.001).

We next quantified factors accounting for the differences in task performance and reaction times. We took advantage of multiple choice target locations used in our task, and measured the dependence of response errors on the distance between the choice targets (Figure 1D i). Successful performance in memory tasks largely depends on two processes: a successful implementation of the task rule, or task bias (Figure 1D ii); and spatially accurate memory representation, or memory tuning (Figure 1D iii).

### Task-rule bias

Earlier studies have reported that failures to implement the appropriate task rule lead to more errors in tasks requiring cognitive control (Antoniades et al., 2013; Klapetek et al., 2016; Munoz & Everling, 2004). Given that choice target locations are selected independently, failure to implement the appropriate task rule would result in an upward shift of the entire tuning function for the Avoid task (Figure 1D i-ii). To investigate this, we compared performance for trials in which choice targets were aligned 180 degrees apart and observed that failures to implement the appropriate task rule increased in the Avoid task (Figure 1E, S3K) (AQ: p<0.001; HB: p<0.001).

### Working memory tuning

The key limiting factor in successful working memory performance is the accuracy of memory representations (Heuer et al., 2020; Ma et al., 2014). In contrast to earlier physiology studies, our task design allowed us to measure spatial memory accuracy in Look and Avoid tasks. If the behavioral strategy of a subject is to memorize the cue location, then there should be no accuracy differences between the two tasks. Alternatively, if the behavioral strategy relies on planning or inhibiting a motor action during the delay period, there should be consistent accuracy differences between the two tasks. Given that the cue location predicts the saccade in the Look task, but not in the Avoid task, memory accuracy is likely to be higher in the former.

We quantified memory accuracy via memory tuning functions, with narrower tuning indicating more spatially accurate memory (Ma et al., 2014) (Figure 1D iii). In a version of the experiment with a high density of response target distances (18 target locations), we found that the Look task exhibited narrower tuning than the Avoid task (Figure 1F, S1A, and S3E) (AQ: p<0.001; HB: p<0.001). We confirmed these results in the main behavioral experiment with 6 target locations (Figure S2C, S3G) and during electrophysiology recording sessions which used 4 target locations (Figure 1G, S3I) (AQ: p<0.001; HB: p=0.003). This result is counterintuitive, as monkeys are highly sensitive to trial outcomes (Platt & Glimcher, 1999) and thus should guide their behavior based on sensory space. Instead, action space has a stronger impact on visual working memory accuracy and reduces successful outcomes in the Avoid task.

We also corroborated this result independent of error trials - a decrease in memory accuracy would likely be associated with increased decision time to select the correct target, thus resulting in longer reaction times (Jonikaitis et al., 2017). Indeed, reaction times also showed tuning differences between Look and Avoid, when response targets were closer together (Figure 1H, S2D, S2F, S3F, S3H, S3I) (AQ 18 targets: p<0.001; 6 targets: p<0.001; 4 targets: p<0.001; HB 18 targets: p<0.001, 6 targets: p<0.001, 4 targets: p>0.05).

Furthermore, we also investigated when the memory accuracy differences between the two tasks were established. In a version of the experiment, we used a wider range of memory period durations (100 to 1200 ms) and observed that memory accuracy differences between Look and Avoid tasks emerged within the shortest measured time - 100-400 ms after the memory cue (Figure 1H and S3J).

Combined, our results show that performance is influenced by a combination of changes in task-rule failures and spatial tuning. This surprising finding reveals that instead of relying on highly accurate sensory space, behavior is strongly influenced by action space.

### Action space reshapes covert and overt attention during working memory

We next measured sensory and action space components in working memory. This is crucial to establish, given lack of such data from earlier neurophysiology studies.

### Covert attention

First, we used a classical reaction time paradigm to measure covert attention (Posner et al., 1980). On a small number of trials (7%), we presented a probe (a single target), and monkeys made an eye movement to it regardless of whether the probe was at the cued or non-cued location (Figure 2A). We presented the probe at different times relative to the cue onset, and saccades to the probe were made as early as 250 ms after the cue onset. Even though probe stimuli were identical in Look and Avoid tasks, reaction times to the probes presented at the cue location were consistently prolonged in the Avoid task, indicating a reduction of covert attentional orienting toward that location (Figure 2B and S5A).

**Figure 2.**
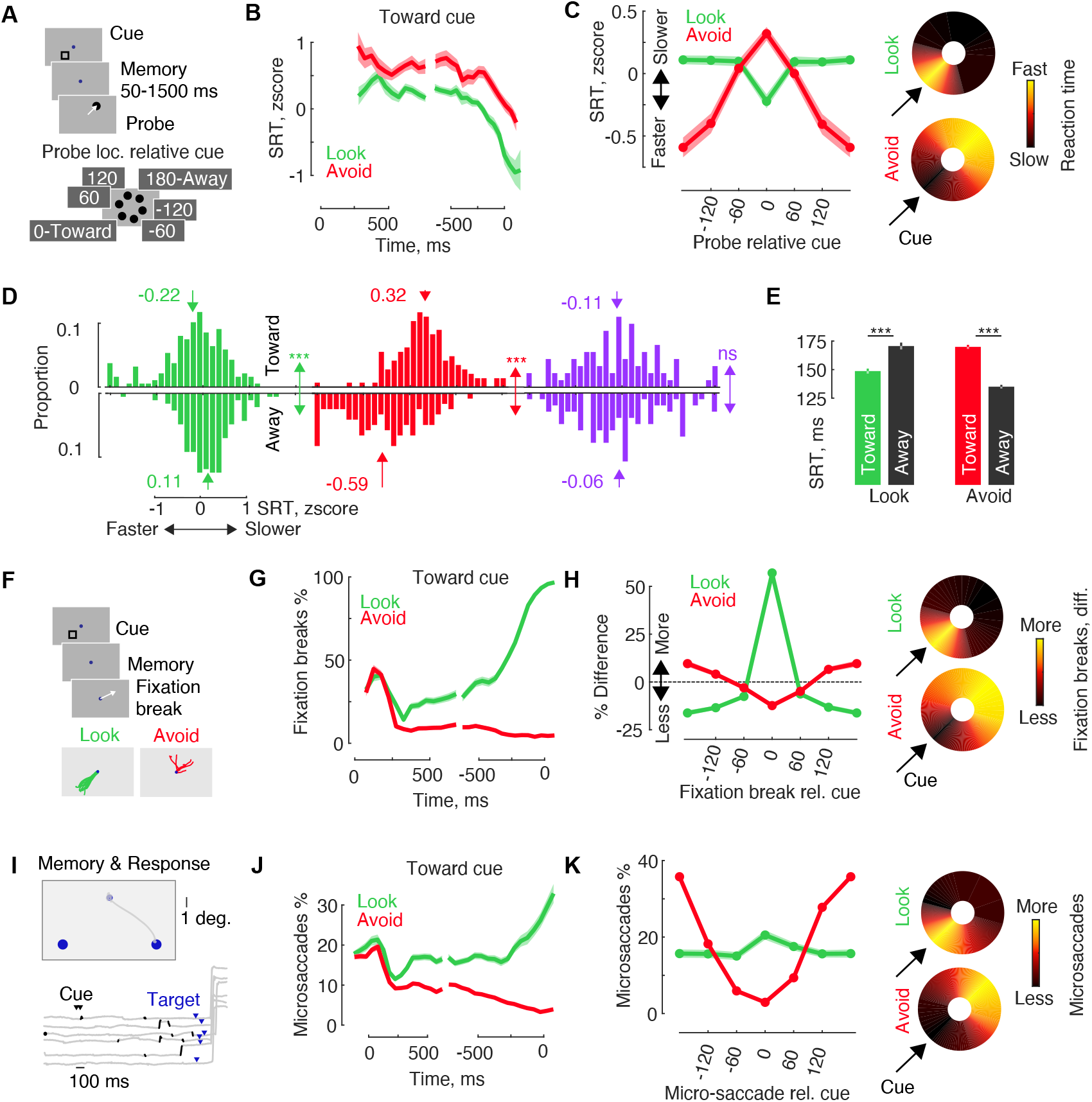
Attentional orienting in Look-Avoid tasks. **(A)** Illustration of a probe trial: Following memory period, a probe appeared at a randomly selected location relative to the cue. “Toward” indicates the probe at the cue location, while “Away” indicates the probe located opposite to the cue. **(B)** Temporal course of saccadic reaction times to the cued location: Time is plotted as the stimulus-response onset asynchrony. **(C)** Saccade reaction times as a function of probe location relative to the cue during the late memory period (last 500 ms of the delay). Heat-maps depict saccadic reaction times, with the slowest and fastest reaction times within each condition defined as the minimum and maximum. Data is rotated so that the cue appears at the lower left, as indicated by the arrow. **(D)** Distribution of individual reaction times toward (upper) and away (lower) from the cue. Significant statistical comparisons between the toward and away conditions are denoted with ***. Purple is Control fixation condition. **(E)** Reaction times during neurophysiology recording sessions when only two probe locations relative to the cue were used. **(F)** Illustration of fixation breaks during the memory period: Insets show eye movement traces during the late memory period in an example session. The data is rotated such that the cue appears at the lower left location. **(G)** Temporal course of fixation break probability toward the cue location. **(H)** Late-Early memory period difference in fixation break probability: Fixation break directions are relative to the cue location. Positive values indicate an increase in fixation breaks toward a location during the late memory period, while negative values indicate a decrease in fixation breaks toward that location. **(I)** Illustration of micro-saccades during the memory period: The upper panel displays an example gaze position during the memory period and choice target periods. Traces below show a few example trials. The cue on-off time is indicated by black arrows, and the saccade target (ST) is indicated by a blue arrow. **(J)** Temporal course of micro-saccades toward the cue location. **(K)** Micro-saccade directions as a function of cue location during the late memory period.

Typically, when sensory and action space overlap, reaction times are faster, indicating covert attentional selection (Carpenter & Williams, 1995; Dorris & Munoz, 1998; Khan et al., 2016; Milstein & Dorris, 2007). We determined the spatial profile of covert attention by presenting the probe at different distances from the cue. Clearly, reaction time profiles were distinct in Look and Avoid tasks - during the late delay period, reaction times were faster to the cue location in the Look task, whereas reaction times to the cue location were slower in the Avoid task (Figure 2C-D, S5B-C) (all p<0.001). Furthermore, in the Fixation control task, which did not require memory of the cue, we observed little to no difference between reaction times to the cue versus non-cued locations (Figure 2D, S5C) (AQ; p>0.05; HB: p=0.02). Lastly, we replicated these findings in neurophysiology recording sessions that used only two probe locations (Figure 2E and S5E) (Look: both monkeys p<0.001; Avoid: both p<0.001). This pattern of data shows that covert attention is aligned with action space, and not with sensory space.

### Broken fixations

On some trials (Figure 2F), monkeys failed to maintain fixation during the memory period. Fixation breaks aborted the trial, provided no reward, and delayed the next trial. First, monkeys broke more fixations in the Look task than in the Avoid task, indicating stronger action preparation during the memory period in the former (Figure S4B, S5E) (AQ: Look-Avoid fixation break difference 9.5±0.81%, p<0.001; HB: 3±0.4%, p<0.001). Furthermore, fixation directions were clearly modulated by the cue location (Figure S4C) and showed distinct time courses. Right after cue presentation, fixation breaks were directed towards the cue in both Look and Avoid tasks (Figure 2G and S5F), indicating oculomotor capture by the visual stimulus (Theeuwes et al., 1998). However, over time, fixation break directions diverged between the Look and Avoid tasks. During the late delay period (Figure S4D, S5H), the proportion of fixation breaks towards the cue was much higher in the Look task (Look-Avoid difference: AQ 84±1.2%, p<0.001; HB 53±3.8%, p<0.001), whereas more fixation breaks were directed away from the cue in the Avoid task (AQ: −31±1%, p<0.001; HB: −18±2.4%, p<0.001).

### Micro-saccades

Micro-saccades are small eye movements observed during periods of fixation (Figure 2I) and are measured on successfully completed trials. Micro-saccades also reveal covert orienting towards attended and memorized locations (Jonikaitis et al., 2019; Vries & Ede, 2023; Yuval-Greenberg et al., 2014). The time-course of micro-saccades (Figure 2J) showed initial, brief orienting towards the visual stimulus, and after that, a diverging probability of orienting between the two tasks. During the late delay, micro-saccade directions were clearly modulated by action space (Figure 2K, S4E, and S5H-I) - more micro-saccades were directed towards the cue in the Look task than in the Avoid task (AQ: 18±1.3%, p<0.001; HB: 2.7±0.6%, p<0.001), and more micro-saccades were directed away from the cue in the Avoid task (AQ: −20±1.3%, p<0.001; HB: −1.5±0.6%, p=0.02).

Furthermore, we observed that the proportion of micro-saccades directed towards the cue on error trials was reduced in the Look task (Figure S4F and S5K, correct versus error difference AQ: 48±10.5%, p<0.001; HB: 14±2.2%, p<0.001). In the Avoid task error trials, orienting towards the cue was increased for one monkey (AQ=-24±2.6%, p<0.001; HB=-1±1.5%, p>0.05).

Combined, using a comprehensive series of behavioral tasks and measures, we observed convergent evidence for action space biasing covert and overt attentional orienting in both the Look and Avoid tasks. This indicates the crucial role action space plays in visual working memory maintenance.

### Sensory versus action space in prefrontal cortex

We next examined sensory and action space-related activity in prefrontal cortex area FEF. We recorded from 4098 single and multi-units across 76 sessions in two monkeys performing the Look and Avoid task. Presentation of the sensory stimulus and maintenance of memory modulated activity in 2084 and 2325 units, respectively (for examples, see Figure S6).

Similar to the behavioral results, we observed that the pattern of neuronal activity during the memory period was different for Look and Avoid tasks. During the visual period, neurons in both tasks showed higher activity for cues presented within the receptive field (Figure 3A, C, and Figure S7A) (cue in vs. cue opposite, p<0.001), confirming sensory space-related responses during the presentation of visual stimuli. During the memory period, however, activity was higher for cues within the receptive field in the Look task (both monkeys combined and individual datasets: all p<0.001) but lower in the Avoid task (all p<0.001).

**Figure 3.**
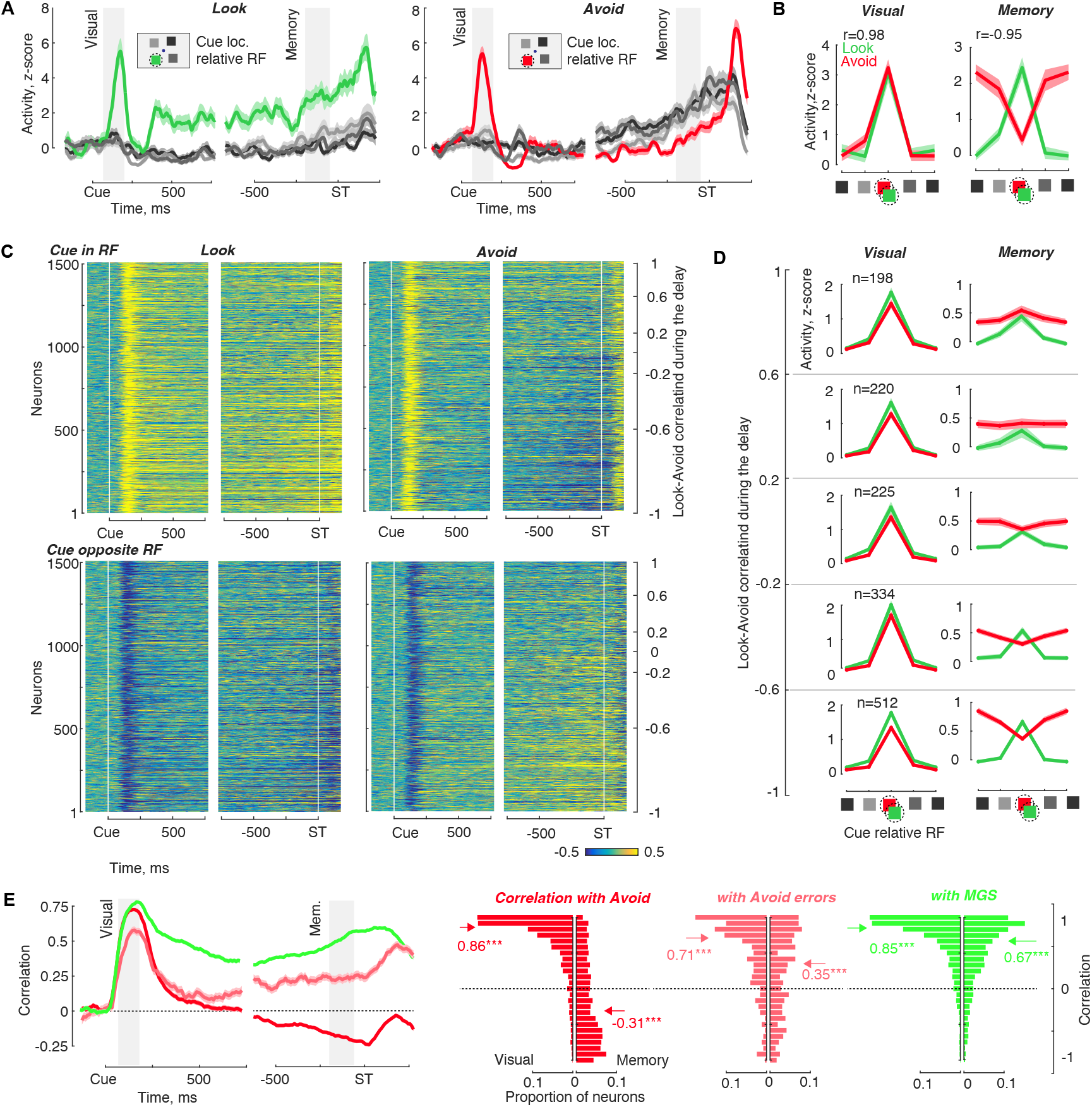
Neuronal activity in Look-Avoid tasks. **(A)** Example neuron recorded in both Look and Avoid tasks. Activity is displayed relative to cue onset (left) and relative to the saccade choice target (ST, right). The legend indicates cue locations relative to the receptive field (dashed circle). Shaded areas indicate the intervals used for data analysis, including the visual period (50 to 200 ms) and delay period (−200 to −50 ms). **(B)** Activity during visual and memory periods (same unit as in A) of Look and Avoid tasks. Activity is presented relative to the center of the visual receptive field. “r” represents the correlation of activity between tuning in Look and Avoid tasks. The unit exhibited a positive correlation during the visual period and a negative correlation during the memory period. **(C)** Heat maps of normalized activity in Look and Avoid tasks. Activity is shown for cues in the visual receptive field (upper) and cues opposite from the visual receptive field (lower) conditions. Neurons are ordered in descending fashion based on tuning correlation between Look and Avoid tasks during the memory period. **(D)** Activity during visual and memory periods of Look and Avoid tasks. Neurons were categorized into groups based on their tuning correlation during the memory period. The top panels represent neurons with a positive correlation, while the bottom panels represent neurons with a negative correlation. **(E)** Tuning correlations between different tasks. The line plot illustrates the between-task tuning correlations over time. Histograms show between-task tuning correlations during the visual (left) and memory (right) periods. Correlations are shown for “Look vs Avoid”, “Look vs MGS”, and “Look vs Avoid error trials”. *** indicates a significant correlation, while “n.s.” represents a non-significant correlation.

We quantified the association of each neuron with sensory or action space by computing spatial tuning correlations between the Look and Avoid tasks (Figure 3B). As expected, positive tuning correlations were dominant during the visual period (Figure 3D-E), signifying a strong resemblance in neuronal responses to sensory stimuli across the two tasks (r=0.86, p<0.001).

However, these tuning correlations diverged during the memory period. In the Look task, neurons maintained a consistent tuning preference between the visual and memory periods, while the majority of neurons in the Avoid task displayed reversed tuning (Figure 3D). As the memory period progressed, the spatial selectivity of neurons during the Look task increased, whereas in the Avoid task, it either decreased or remained unchanged (Figure S7D). Taken together, an anti-correlation between Look-Avoid tuning emerged during the memory period (Figure 3E) (r=-0.31, p<0.001). This implies that neurons with the highest memory-related responses in the Look task also demonstrated the most pronounced inverted responses in the Avoid task.

Moreover, error trials of the Avoid task were positively correlated with the Look task (Figure 3E) (r=0.35, p<0.001), indicating that neuronal responses during the memory period were linked to behavioral responses. Additionally, as a control comparison, we found high positive correlations between Look and MGS trials during both the visual (r=0.85, p<0.001) and memory periods (Figure 3E) (r=0.67, p<0.001).

FEF neurons exhibit diverse responses, with some being most responsive to the visual, memory, or motor periods in working memory tasks, respectively (Bruce & Goldberg, 1985). We investigated whether the reduced correlation in Look-Avoid activity was equally observed in sub-populations of neurons that were most responsive to different task epochs (Figure S7E). All sub-populations with memory period activity displayed anti-correlation. However, the strongest decrease in correlation was observed in populations with motor activity (“Memory and Motor”: r=-0.49, p<0.001; “Visual, Memory, and Motor”: r=-0.47, p<0.001), while a weaker decrease was observed in populations without motor activity (“Visual and Memory”: r=-0.21, p<0.001). The other sub-populations showed no correlation between Look and Avoid (“Visual only”: r=-0.01, p=0.15; “Memory only”: r=0.001, p=0.75; not enough “Motor only” units for this analysis). These findings further support the hypothesis that memory period signals are related to action, rather than sensory space.

### Linear and mixed selectivity during target choice

We next tested whether memory period activity is associated with behavioral choices. We found that during the target period - after the memory period and before monkeys made an eye movement - neuronal responses to the target at the saccade location were always higher (Figure 4A; Look p<0.001, MGS p<0.001, Avoid p<0.002, Avoid errors p<0.001), indicating linear neuronal selectivity across different tasks during target choice (Rigotti et al., 2013).

**Figure 4.**
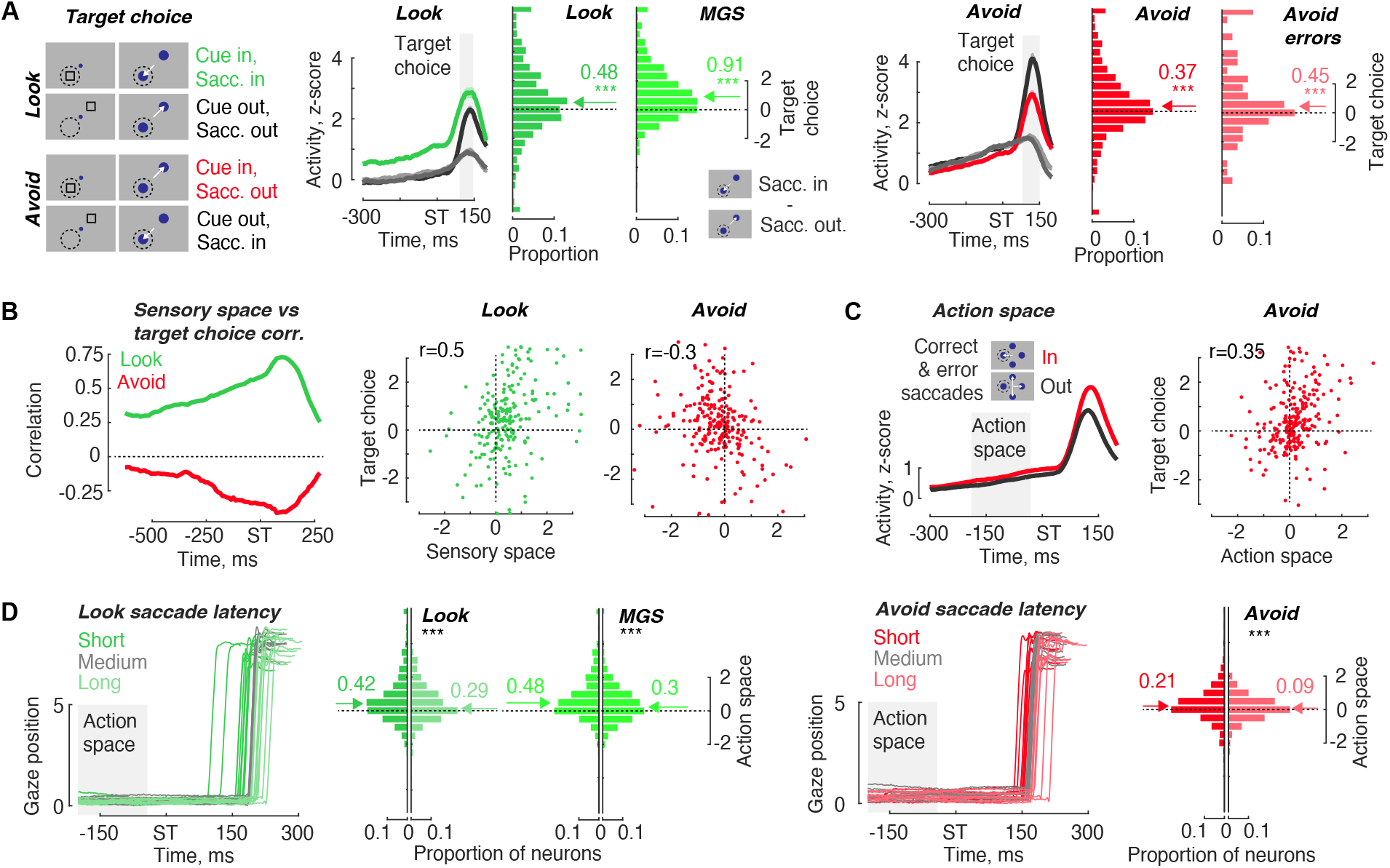
Action space and target choice modulation. **(A)** Schematic illustration of target choice. Target choice is defined as the activity difference between “saccade in RF” and “saccade opposite RF”. The time course of target choice in Look and Avoid tasks is represented as line plots. The shaded area indicates the target period used for analysis (100 to 150 ms, trials with shorter saccade latencies are excluded). Histograms display the distribution of target choice for the neuronal population. **(B)** Within-task correlation between sensory space and target choice. Sensory space is defined as the activity difference between “cue in RF” and “cue opposite RF”. Sensory space at each time point is then correlated with target choice (100 to 150 ms). Scatter plots visualize the relationship between sensory space during late delay period (−200 to −50 ms) and target choice in Look and Avoid tasks. “r” represents the correlation coefficient. **(C)** Action space in Avoid task. Action space is defined as the difference between “saccade in” versus “saccade out” trials, encompassing correct and error trials. Crucially, action space is defined during the memory period (−200 to −50 ms), when the action might be spontaneously suppressed or anticipated, but the correct action location is not yet known. Scatter plots depict the relationship between action space and target choice in Look and Avoid tasks**. (D)** Action space during long and short latency trials. Line plots illustrate an example session divided into short (fastest 33% of saccades) and long (slowest 33% of saccades) saccade latencies. Action space was then calculated for each neuron separately for short and long latency trials. *** indicates a significant difference in action space between short and long latency trials.

We then calculated correlations between target choice (saccade in vs. saccade out difference during the target period) and sensory space (cue in vs. cue out difference during the memory period). Correlations were positive in the Look task (Figure 4B, r=0.5, p<0.001), negative in the Avoid task (Avoid r=-0.30, p<0.001), and positive again in the Avoid error trials (r=0.11, p<0.05), indicating mixed neuronal selectivity between target choice and sensory space. However, in the Avoid task, we recalculated the correlation between target choice and the action space. Crucially, action space is defined during the memory period, when the action might be spontaneously suppressed or anticipated, but the correct action location is not yet known (future saccade in vs. saccade out difference during the memory period). We observed a positive correlation (Figure 4C) (Avoid r=0.35, p<0.001), showing linear selectivity.

We further tested whether the action space influenced target choice by dividing each experimental session into trials with short and long eye movement reaction times. In the Look, MGS, and Avoid tasks, short eye movement reaction times were associated with higher action space-related activity (Figure 4D) (Look p<0.001, MGS p<0.001, Avoid p<0.001). Combined, the results show the crucial role of delay activity in determining behavioral choices.

### Memory accuracy differences in neuronal responses

We next investigated whether neuronal activity could be associated with the memory accuracy differences between the Look and Avoid tasks. For this purpose, we utilized linear decoders (SVM) to decode memory location from the activity of concurrently recorded neuronal populations. We built the decoder for each session (simultaneously recorded neurons) and averaged the performance across sessions. We observed a rapid increase in decoding accuracy after cue onset in both tasks (Figure 5A, Figure S10A). During the memory period, the decoding accuracy remained robustly above chance level in both tasks, yet it was higher in the Look task (Figure 5A, S10B) (p<0.001). Furthermore, the decoder weights were positively correlated after cue onset between Look and Avoid task (Figure S10B) (r=0.59, p<0.001), but were negatively correlated during the memory period (Figure 5B) (r=-0.18, p<0.001). This suggests that independent decoders in the Look and Avoid tasks utilized an overlapping set of neurons, but with reversed weights, consistent with the correlation results.

**Figure 5.**
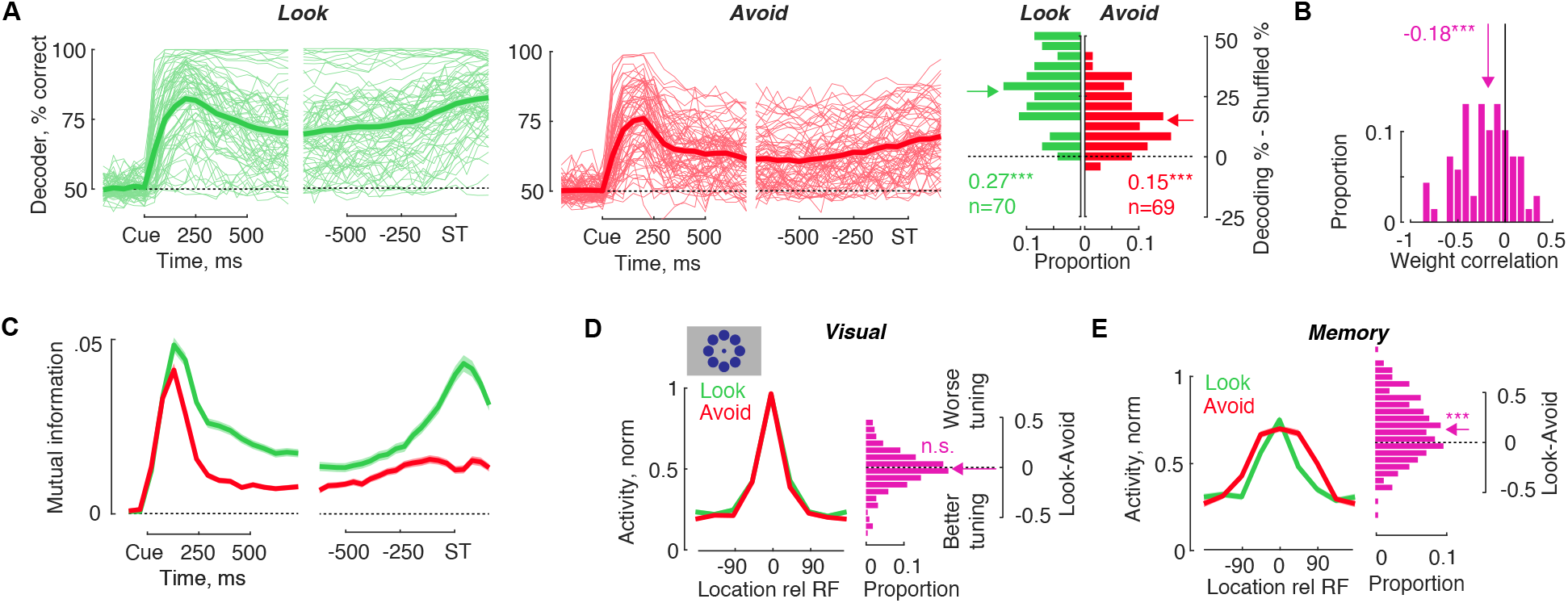
Memory accuracy differences in neuronal responses. **(A)** Decoding cue versus opposite location in Look and Avoid tasks. Decoding was performed for each individual session using simultaneously recorded neuronal populations. Thin lines represent individual sessions, while the thick line represents the average across all sessions (N=70 for Look task and N=69 for Avoid task). The histogram displays the distribution of decoder accuracies during the memory period, minus shuffled label decoding. Positive values indicate that the decoding performance is higher than chance level. **(B)** Distribution of Look-Avoid task correlations of decoder weights. **(C)** Time course of mutual information in Look and Avoid tasks. **(D)** Spatial tuning in a version of the experiment with 8 target distances (number of neurons = 212). Neuronal activity is normalized to a range between 0 to 1 for each task. Better or worse tuning is determined as the activity difference between Look and Avoid tasks at locations 60-120 degrees away from the receptive field. Tuning differences are measured during both visual and memory periods.

We also computed mutual information, which captures both how much information about the memory cue location is conveyed by the neuronal activity and the reliability of this information (Figure S10D). After the cue presentation, mutual information increased in both tasks, but rapidly differentiated between the two (Figure 5C). By the end of the memory period, mutual information was higher in the Look task (Figure S10E) (p<0.001).

In a follow-up version of the experiment with more memory locations (8 locations), we measured the neuronal tuning in both the Look and Avoid tasks (n=237). To make a comparison between tuning curves, we normalized the neuronal responses to the maximum response and shifted the tuning functions in the Avoid task to match the peak with the Look task. During the visual period, neurons in both tasks showed similar tuning (Figure 5D) (p>0.05). However, during the delay period, we observed that neuronal responses were more spatially tuned in the Look task than in the Avoid task (Figure 5E) (p<0.001). Taken together, our findings demonstrate that the neuronal activity maintains more information about the cue location in the Look task, in line with the behavioral results.

### Stability and transformation of working memory

A reversal of correlation from the visual to memory period indicates a transformation from sensory to action space in working memory. We employed a cross-temporal linear classifier (SVM) to test whether the neuronal activity was associated with stable or dynamic coding of memorized stimuli (Murray et al., 2017; Spaak et al., 2017). The linear classifier was constructed by training at one time point (t) and testing at different time points (t+delta) (Figure 6A). If the decoding was robust only when the test and training intervals overlapped, this indicated a dynamic pattern. Alternatively, if the decoding accuracy generalized between two distinct and temporally displaced time intervals, it was indicative of a stable pattern.

**Figure 6.**
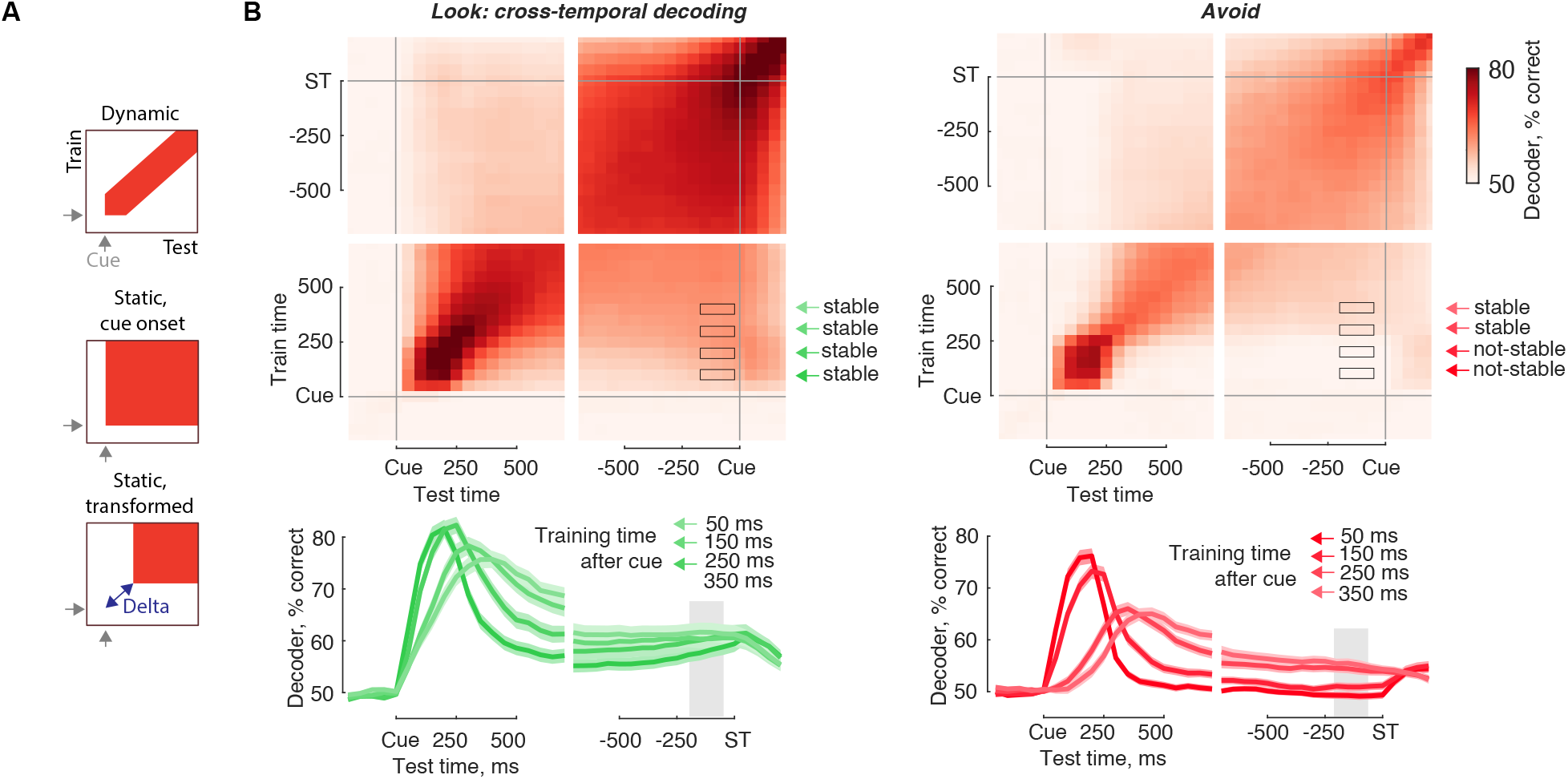
Sensory to action space transformation. **(A)** Illustration of dynamic and stable population responses. Successful decoding along the diagonal indicates a dynamic population response. Successful decoding in the area marked with a square indicates a static population response. A static population response can begin immediately after cue onset or be delayed in time by a value labeled as “Delta”. **(B)** Cross-temporal decoding in Look and Avoid tasks. A linear decoder was trained and tested on simultaneously recorded neuronal populations at different times during the trial. Line plots display decoders trained at four different times after the cue (50, 150, 250, 350 ms) and tested throughout the trial. For each of these four training times, memory representation was labeled as stable if decoder performance was above chance at the end of the memory period. The time interval at the end of the memory period is marked with black frames on the heat map and shaded area on the line plot.

In the Look task, the cross-temporal population pattern exhibited evidence of stable coding (Figure 6B). This stable coding emerged after the onset of the visual cue, indicating that sensory space and action space overlapped throughout the delay period. Decoding successfully generalized to the end of the delay period for the first time bin after the cue (Figure 6B) (50 ms, p<0.001), suggesting that prefrontal activity can bridge the time interval from sensory stimulus to action response. We also validated this result using cross-temporal tuning correlations (Figure S11A-B) (50 ms, p<0.001).

In the Avoid task, we identified a stable cross-temporal population pattern as well; however, it occurred with a delay (delta, as shown in Figure 6A) - it took 200-250 ms following cue onset to emerge (Figure 6B) (p<0.001). This timing was further confirmed by using neuronal tuning correlations (Figure S11B). Our results strongly suggest that population activity evoked by visual input is transformed in the Avoid task, supporting the hypothesis that sensory space is transformed to action space during the memory period.

To explore whether sensory and action space can be generalized across two tasks, we trained a decoder at one point (t) in the Look task and tested it at a different time point (t+delta) in the Avoid task. Surprisingly, our results showed that memory period activity in the Look task could successfully decode visual activity in the Avoid task (Figure S11C) (p<0.001), demonstrating a generalization of sensory space across two tasks. However, memory activity in the Look task was associated with small, but significant below-chance decoding accuracy of memory activity in the Avoid task (p=0.01), highlighting the role of action space in the population activity.

In summary, our findings establish the time course of visual input manipulation in working memory, revealing that manipulation occurs immediately or in parallel with visual information presentation, and takes 200-250 ms to accomplish.

## Discussion

The function of working memory is to store past sensory inputs, yet the goal for this storage is to guide future behavior. However, physiological mechanisms of working memory are typically investigated as if memory was independent from future actions. Here we used two tasks in which sensory information to be maintained in working memory was identical, but future actions differed. The Look task relied on direct stimulus-action mapping during working memory, whereas the Avoid task transformed this mapping.

### Action space

Our findings provide compelling behavioral and physiological evidence for the role of action space in shaping working memory representations (Ehrlich & Murray, 2022; Schneider, 1999). We observed that working memory representations rapidly diverged between Look and Avoid tasks. Behaviorally, this was evident as a reduction of spatial tuning for the Avoid task, whereas a decrease in mutual information and in memory decoding was observed in neuronal activity. This indicates that mere anticipation of different actions can strongly bias working memory. Previously, it has been hypothesized that action anticipation could protect working memory from distraction or reduce representational complexity (Henderson et al., 2022; Myers et al., 2017). In line with this, our findings suggest that action preparation towards the cue location in the Look task is associated with better memory accuracy, and suggest that performance in Look or MGS tasks is likely a function of spatial memory and action preparation.

Earlier studies frequently relied on dual-task paradigms, in which participants memorized sensory stimuli and then completed an intervening eye or hand movement during the working memory period. These eye or hand movements improve working memory if they are directed to the memorized stimulus and impair if directed away (Hanning et al., 2016; Lawrence et al., 2001; Ohl & Rolfs, 2018, 2020). Instead of relying on dual-task context, here we employed a novel approach by manipulating the relationship between working memory and motor response. This approach leaves sensory space intact; it does not add dual-task burden, but it alters the relationship between sensory stimulus and the action. Our findings demonstrate that this is an effective way to investigate the relationship between working memory and representation space.

### Attention-memory-saccades

It has been proposed that there is a significant overlap between working memory, visual attention, and action preparation (Awh et al., 2006; Jonikaitis & Moore, 2019). Here we show that action space plays a major role in determining this overlap. In the Look task, these three cognitive processes are centered on the same location, and as expected, we observed action preparation and attentional orienting towards the sensory memory location (Binda & Murray, 2015; Jonikaitis et al., 2019; Mathôt et al., 2013; Yuval-Greenberg et al., 2014). Avoid task, on the other hand, poses a conflict - attending to a memorized location can lead to an involuntary oculomotor capture (Theeuwes et al., 1998), which would decrease task performance. We observed such capture during the early memory period, but it rapidly decreased during the Avoid task (see also Dhawan et al., 2013; Jonikaitis et al, 2019).

In line with behavior, neuronal activity was also modulated by action space, with most neuronal responses higher at cued locations in the Look task, and lower at cued locations in the Avoid task. Combined, these findings support the hypothesis that action space determines the location of attention and spatial working memory (Baldauf & Deubel, 2010; Rizzolatti et al., 1987). Our findings show that accounting for action space is crucial when investigating neuronal activity associated with working memory.

### Relationship to other tasks

Several different tasks have been used to investigate spatial working memory, including delayed response, sequence memory, and dissociated stimulus-response tasks. In delayed response and sequence memory tasks, sensory stimuli or multiple stimuli are also response targets (Funahashi et al., 1989; Xie et al., 2022). Indeed, it has been shown that responses are prepared for single targets or multiple targets simultaneously (Baldauf & Deubel, 2008a; Godijn & Theeuwes, 2003; Jonikaitis et al., 2017), indicating a substantial role that action preparation could play in observed memory period activity. In dissociated stimulus-response tasks, the visual stimulus predicts a spatially displaced response, such as an action target rotated 90 or 180 degrees away from the sensory stimulus (Funahashi et al., 1993; Munoz & Everling, 2004; Takeda & Funahashi, 2002). However, research has also indicated that multiple motor actions compete in anti-saccade tasks, leaving the question open as to whether a mixture of sensory stimulus and action preparation determines memory period activity in such tasks. Spontaneous behavioral strategies formed during the Avoid task, on the other hand, can clearly dissociate sensory and action components of working memory, suggesting that this task is uniquely suitable for investigating action space.

### Cognitive control

The Avoid task, by altering direct stimulus-action associations, can be considered a cognitive control task (Aron, 2011; Cai et al., 2011; Verbruggen et al., 2019). Earlier studies of cognitive control have suggested that action suppression plays a key role in tasks that modify reflexive stimulus-action sequences (Aron, 2011; Noorani & Carpenter, 2013; Verbruggen & Logan, 2008). This view provides a potential insight into understanding the data in the Look and Avoid tasks. In the Avoid task, we observed initial orienting towards the sensory memory location, followed by a reduction in orienting to it. This could mean that action space might involve the suppression of the sensory stimulus location and, therefore, a relative facilitation of locations away from the sensory stimulus (this also indicates that “action space” describes diverse results better than “action preparation”). Suppression of actions towards a sensory stimulus could be a likely strategy instead of trying to predict multiple potential response target locations in the Avoid task. Indeed, we varied the number of potential targets in different versions of the experiment, and memory tuning was comparable between these different versions. Suppression of actions to a sensory stimulus location could thus produce memory tuning that is different from the facilitation observed in the Look task (Jonikaitis et al, 2019). However, our findings also highlight that disentangling action selection and suppression components of cognitive control might be much harder than previously assumed. This is due to the fact that the selection of the cued location in the Look task was simultaneously associated with reduced orienting away from the cue, and suppression in the Avoid task was associated with a parallel increase in orienting away from the cue. Similarly, neuronal activity was also simultaneously increased and reduced at different locations relative to the cue in the Look and Avoid tasks.

### Transformations in working memory

Our findings build upon earlier work on dynamic and stable patterns of activity in neuronal populations in the prefrontal cortex (Murray et al., 2017; Spaak et al., 2017). Within the Look task, the population response to the sensory stimulus generalized to the decoding of activity during the memory period. This rapid formation of stable population activity has also been suggested by a recent study with human participants, showing that motor coding is activated whenever the sensory stimulus allows for sensory to action mapping (Ede, Chekroud, Stokes, et al., 2019). Furthermore, Look memory period activity generalized to Avoid sensory responses, establishing stable population patterns of activity within and between tasks.

In the Avoid task, stable population patterns were formed only after sensory memory transformation to action space. This manipulation and transformation are key properties of working memory (Baddeley, 1992), and we show that this process can be achieved as rapidly as 200 ms. This transformation resulted in a shift of neuronal tuning away from the sensory stimulus location and a reduction of covert and overt behavioral orienting towards the sensory memory. Here, we demonstrate that this manipulation was accomplished within working memory, independent of the sensory stimulus availability. Earlier studies typically focused on manipulations of sensory stimuli, using sensory stimuli durations that are much longer (500-1000 ms) than the time it took to accomplish this manipulation in the current study. Therefore, we show that this manipulation can be achieved in working memory without reliance on the sensory stimulus duration. We further observed that this manipulation was crucial for behavior, as successful transformation in neuronal tuning was associated with faster behavioral responses and correct-error responses.

### Role of FEF in working memory

Findings from earlier studies allow for the possibility that neuronal activity could be correlated to sensory space in other areas, such as dlPFC or LIP (Funahashi et al., 1989; Pesaran et al., 2002; Kojima, 1980; Spaak et al., 2017; Wasmuht et al., 2018; Zhang & Barash, 2004). However, earlier studies did not evaluate contributions of sensory space and action space to neuronal activity in those areas. Our results suggest that there are multiple open questions with respect to neuronal working memory signals. First, our findings open the possibility that action space could have influenced neuronal activity observed in other visual memory studies. Here, we provide a behavioral template on how such influence can be measured. Second, if sensory space is maintained in other cortical areas, the contribution of sensory space to influencing behavioral choice remains to be demonstrated. Future research should focus on investigating how sensory space is utilized and which brain structures are involved in the storage of sensory and action space. Our findings suggest that action space has a major impact on behavioral actions and neuronal activity in FEF.

In summary, our study provides compelling evidence that working memory is highly influenced by the anticipation of future behavioral actions. The conventional view of working memory as the simple storage of past sensory inputs needs to be reconsidered in light of the crucial role of action space. Overall, our study opens up new avenues for exploring the complex interplay between working memory and action.

## Methods

Two male rhesus macaques (Macaca mulatta), weighing 11 kg and 14 kg, were used in this study, named Aquaman (AQ) and Hellboy (HB) respectively. All experimental procedures adhered to the ethical guidelines outlined in the National Institutes of Health Guide for the Care and Use of Laboratory Animals, the Society for Neuroscience Guidelines and Policies, and were conducted under the approval of the Stanford University Animal Care and Use Committee (IACUC) protocol (#APLAC-9900). Further details of the experimental procedures can be found in a previous report (Armstrong et al., 2009).

### Behavioral tasks

Each behavioral trial commenced with a blue circle fixation spot (radius of 0.5° visual angle; luminance: 3.8 cd/m2, RGB color: 0.08, 0.08, 0.78) displayed at the center of a gray background screen (luminance: 10.7 cd/m2, RGB color: 0.5, 0.5, 0.5). Once the monkey acquired fixation (within 0-200 ms), a task-irrelevant texture was presented on the screen and remained visible throughout the trial (details below). After maintaining fixation for 600-800 ms (duration chosen randomly for each trial), a cue appeared as a colored square frame (size of 1° x 1° visual angle) for approximately 50 ms at a randomly selected location (1 location selected from 4-18 possible locations; eccentricity from fixation ranged from 5° to 7° visual angle across different sessions; location selected randomly for each trial, independent of the previous trial). Following cue presentation, a delay period ensued (duration selected randomly for each trial) lasting 1400-1600 ms (with a subset of sessions having delay periods as short as 200 ms or as long as 2000 ms). After the delay period, the fixation spot disappeared, and one of four behavioral response options was presented (details below). Monkeys received a juice reward for making a correct saccadic eye movement and maintaining gaze on the target for 200 ms. The inter-trial duration after a correct response was 100 ms. Failures to acquire fixation, breaks of fixation during the trial, or incorrect eye movements were not rewarded and were followed by a 2000 ms inter-trial duration. All stimuli and task parameters for each new trial were randomly selected and independent of the previous trial to prevent task switching costs. To reduce typically observed cognitive task-switching costs (Antoniades et al., 2013), different behavioral tasks (Look-MGS, Avoid, and Fixation control) were conducted in separate blocks, with the block duration varying from 150 to 400 trials based on the monkey’s motivation. Each session could start with either a Look-MGS or an Avoid block.

### Look-MGS task block

For monkey AQ, the cue color was black and 2 represented by an open square (luminance: 0.2 cd/m^2^ , RGB color: 0.08, 0.08, 0.08), while for monkey HB, the cue color was green and represented by an open square 2 (luminance: 20.1 cd/m^2^ , RGB color: 0.08, 0.78, 0.08).

Look and MGS task trials were randomly interleaved. On Look trials (approximately 46.5% of trials), after the delay period, fixation disappeared, and two targets appeared as filled blue circles (radius of 1° visual angle, luminance: 3.8 cd/m^2^, RGB color: 0.08, 0.08, 0.78). One target always appeared at the previously cued location, while the other appeared at a randomly selected location among the remaining ones (location selected randomly for each trial). To receive a reward, monkeys had to make a saccadic eye movement to the target at the cued location.

On MGS trials (approximately 46.5% of trials), after fixation disappeared (and no targets appeared), monkeys were required to make an eye movement to the memorized location of the cue. If the saccade was directed within a 5° visual angle from the cued location, a target (filled blue circle) appeared to confirm the correct response location, and a reward was provided.

### Avoid task block

In Avoid trials, the setup was identical to Look trials, with the only difference being the rewarded saccadic eye movement. In Avoid trials, monkeys were required to make a saccadic eye movement to the novel target, which was the one not previously cued. The cue color indicated the task, with green cue color for monkey AQ and black cue color for monkey HB.

### Fixation control task block

During fixation control trials, the cue color was white, represented by a filled square (RGB color: 1, 1, 1). In these trials, monkeys were required to maintain their gaze at the central fixation spot until the end of the delay period. To receive a reward, no response saccadic eye movement should occur during the trial.

### Probe trials

On approximately 7% of trials, randomly interleaved within either Look-MGS, Avoid, and Fixation control blocks, after the delay period, only one target appeared. The target was represented by a filled black circle (RGB color: 0.08, 0.08, 0.08). The target could appear at either the cued or non-cued locations, and monkeys had to make a saccadic eye movement to the probe target to be rewarded.

### Cue locations and delay durations

The number of cue locations varied based on the goal of the experiment, as depicted in Figure S1A. For behavioral recording sessions, 6 cue locations were used, spaced 60° apart in polar angle. In behavioral sessions for the memory accuracy experiment, 18 target locations were used, spaced 20° apart in polar angle. For neurophysiology experiments, 4 cue locations were used, spaced 90° apart in polar angle, to ensure sufficient numbers of trials for neuronal activity. In a small subset of neurophysiology experiments for memory tuning, 8 locations were used (across 11 recording sessions).

For a typical behavioral or neurophysiology recording session, memory period durations were 1400-1600 ms. However, in the memory tuning over time experiment, memory period durations were randomly selected from the range of 200-1200 ms. On some sessions, memory durations could slightly vary, based on monkey motivation, but they were always randomly selected.

### Background texture

The screen background was filled with a task-irrelevant texture starting 600 ms before cue onset (0-200 ms after the monkey acquired fixation) (Supèr et al., 2001). The background texture consisted of a dense field of 10,000 uniformly oriented lines (width: 2 pixels, length: 2° visual angle, RGB color: 0.36, 0.36, 0.36). Each background had one orientation selected randomly from 0° to 179° in 30° increments.

On approximately 5/6 of the trials, the background texture was presented, while on approximately 1/6 of the trials, no texture was presented, and the background remained a uniform gray. The probability of having no texture or each texture angle was thus approximately 1/6.

Halfway through the delay period, a new texture or no-texture background was presented, selected randomly and independently of the first texture. Texture presentation was used to evoke visual responses during recordings in visual area V4, and matching parameters were maintained in FEF recordings. It’s worth noting that FEF neurons were typically not responsive to either texture onset or texture orientation.

### Behavior and neurophysiological recording procedures

Experiments were controlled by a DELL Precision Tower 3620 desktop computer and implemented in Matlab (MathWorks, Natick, MA, USA) using Psyctoolbox and Eyelink toolboxes (Brainard, 1997; Cornelissen et al., 2002). Eye position was recorded with an SR Research EyeLink 1000 desktop-mounted eye-tracker for online gaze position tracking (sampling rate 60 Hz) and for offline analysis (sampling rate of 1000 Hz). Stimuli were presented at a viewing distance of 60 cm on a VIEWPixx3D display (1920 x 1080 pixels, vertical refresh rate of 60 Hz).

Neuronal recordings were obtained using 16, 24, and 32 channel linear array electrodes (based on the availability of the electrodes) with contacts spaced 75 or 150 μm apart (U-Probes, V-Probes, and S-Probes, Plexon, Inc). The electrodes were lowered into the cortex using a hydraulic microdrive (Narishige International) at angles roughly perpendicular to the cortical surface. Neuronal activity was measured against a local reference, a stainless guide tube, which was close to the electrode contacts. Plexon data were amplified and recorded using the Omniplex system (Plexon Inc., Dallas, TX). Wide-band data filtered only in hardware at 0.5 Hz highpass and 8 kHz lowpass were recorded to disk at 40 kHz. Some recordings (n=12) were obtained using Neuropixels probes (Neuropixels 1.0 NHP short or long probe, IMEC Inc, Belgium), which contain 384 electrode contacts that could be simultaneously selected for recording. Every 2 contacts are arranged in one row, and the vertical space between rows is 20 μm. Data collection was performed using SpikeGLX software and sampled at 30 kHz. Raw wide-band data were pre-processed with median subtraction and high-pass filtered at 150 Hz.

For both Plexon and Neuropixels recordings, spike sorting was performed using Kilosort2 software (Pachitariu et al., 2023) (https://github.com/MouseLand/Kilosort) and manually curated with Phy (https://github.com/cortex-lab/phy) to remove atypical waveforms and perform minimal merging and splitting. Double-counted units were removed according to previously reported criteria (Siegle et al., 2021). Some key parameters in Kilosort2 we used were: Ops.th=[10,6]; Ops.lam=20; Ops.AUCsplit=0.8; Ops.ThPre=8; Ops.spkTh=-6.

The FEF was localized based on its neurophysiological characteristics and the ability to evoke saccades with electrical stimulation. Electrical microstimulation consisted of 100-ms trains of biphasic current pulses (0.25 ms, 200 Hz) delivered with a Grass stimulator (S88) and two Grass stimulation isolation units (PSIU-6) (Grass Instruments). The FEF was defined as the region from which saccades could be evoked with currents <50 μA (Bruce et al., 1985). After successful localization of FEF, the follow-up recordings were then completed in the area surrounding the micro-stimulated site.

Visual receptive fields were also mapped by presenting briefly shown visual stimuli at different locations on the screen (Figure S1C). Neuronal responses to the onset of visual stimuli were processed online and visualized as visual response maps. These response maps were used to infer locations of visual receptive fields and then to position the memory cue relative to the location evoking responses in most of the neurons.

### Eye movement analysis

Online gaze position was used to update trials and provide appropriate reward feedback. Gaze position was also recorded for offline data analysis. Gaze position on each trial was offline drift-corrected using the median gaze position from the 10 previous trials. Drift correction was based on gaze position from 100 ms to 10 ms before the cue onset when stable fixation was maintained (trials with saccades larger than 1 degree were excluded from median gaze position calculations). We detected saccades offline using an algorithm based on eye velocity changes (Engbert & Kliegl, 2003). We then clustered saccades as ending on one of the three potential locations: (1) fixation, (2) correct response target, (3) wrong response target. The clustering procedure used a support vector machine algorithm with a Gaussian kernel (Donatas Jonikaitis et al., 2019). Saccades directed to the target or distractor had a latency of at least 50 ms after the response cue (Fischer & Boch, 1983), and saccades occurring at shorter latency or during the memory period were classified as fixation breaks. For the microsaccade analysis, we used all saccades that did not break the fixation window during the pretrial period - typically saccades with amplitudes less than 1° visual angle. We removed trials if blinks occurred from 100 ms before cue onset to 200 ms after the time of saccade target onset. Data from each recording were inspected for saccade detection accuracy and data recording noise.

#### Behavioral data analysis

We used correct trials (correct target selected), unless specified otherwise. Responses to the target in the Look-MGS task were separated for Look and MGS trials, as Look responses are typically faster than MGS responses (Figure S2B, S4D). Given that fixation breaks and microsaccade directions are continuous polar angle variables, the data was binned into 6 bins centered on the 6 cue locations used in the behavioral tasks.

#### Statistical comparisons

For statistical comparisons of paired means, we drew (with replacement) 10000 bootstrap samples from the original pair of compared values. We then calculated the difference of these bootstrapped samples and derived two-tailed p-values from the distribution of these differences. For repeated measures analyses with multiple levels of comparisons, we used one-way and two-way repeated measures ANOVAs. All correlations were computed as Pearson coefficients. For behavioral data, one data point represented one session. For neurophysiological data, one data point represented a single or multi-unit neuronal response.

#### Neuronal data curation

To ensure the stability of the recorded data’s firing rate, we identified specific time intervals where significant changes in the firing rate occurred. For each neuron, we calculated the average spiking rates for every trial (with each trial representing one data point). Next, we divided this data into 2-4 continuous trial blocks using the Jenks Natural Breaks detection algorithm (specifically, the “get_jenks_interface” function from the Matlab exchange). The Jenks Natural Breaks algorithm is a clustering method that effectively detects abrupt changes in noisy data. It helps identify the index within the data where high and low values are separated, which can indicate instances when a neuron stops responding during a recording session. By applying this algorithm, we determined whether a neuron either stopped responding (if the spiking rate fell below 0.3 spikes/second for more than 50 consecutive trials) or experienced a drastic decrease in neuronal activity (if the spiking rate fell below 10% of the median spiking rate in the experiment for more than 50 consecutive trials). If any of these conditions were met, the corresponding trial blocks were excluded from the data analysis. Additionally, if a neuron was active in less than 20% of the experimental trials, it was completely removed from further analysis.

#### Data normalization

For each trial, we normalized spike rates by computing a z-score of each neuron’s firing rate on each each trial using a sliding window of ten trials before and after the current trial (Wasmuht et al., 2018): (FR(k,t,j) - mu(t0,j)) /sigma(t0,j)

FR(k,t,j) is the firing rate at trial k, time-bin t, for neuron j; mu(t0, j) and sigma(t0, j) are the mean and standard deviation of the neuron j firing rate estimated from the 21 trials centered on k, for pre-cue time interval t0 (−300 ms to 0 ms before cue onset). Given that mu and sigma are centered on k, first 10 trials of each recording used mu and sigma centered on 11th trial; last 10 trials used mu and sigma centered on 11th trial from the end of the recording.

#### Visual, memory and motor activity

The visual, memory, and motor FEF units were classified using the Look-MGS and Avoid tasks. Motor units were defined using only MGS trials (no visual stimulus was presented at the saccade target location before the saccade on those trials). A unit was classified as a visual or memory unit if its activity was modulated by different cue locations, as estimated by a significant ANOVA main effect (p<=0.05) for either the Look-MGS or Avoid task (Hasegawa et al., 2004). Visual units included activity from 50 to 200 ms after cue onset. Memory units included activity from −200 to −50 ms before the memory period ended. Motor units were classified and included activity from 150 to 0 ms before saccade onset.

#### Neuronal response modulations

We employed two approaches to measure neuronal responses to cue/target locations. The first approach involved calculating the difference in responses between the stimulus in the receptive field (RF) and the opposite RF, which provided a measure of spatial selectivity during the visual or memory period. This measure relies on the detection of the visual RF within a window of 50-200 ms after cue onset and is commonly used to determine “spatial modulation.”

In contrast, the analysis of tuning correlations does not rely on the detection of visual RF. This analysis is equally applicable to neurons with visual actibity, neurons with memory activity but no visual activity, as well as neurons with slower visual responses (than the 50-200 ms window). In this analysis, the mean responses of each neuron to cue locations (4 or 8 different locations based on neurophysiology recordings) in one condition were correlated with responses in another condition (such as Look versus Avoid or cue period versus memory period). The resulting tuning correlations range from −1 to 1. Positive correlations indicate linear selectivity, implying similar tuning between two tasks or time points, while negative correlations indicate mixed selectivity with divergent tuning.

In comparisons involving neurophysiology sessions with 8 targets, the data was normalized within the range of [0 to 1]. During the cue period, the data was rotated based on the cue location relative to the receptive field, with 0 corresponding to the cue at the receptive field. During the memory period, the data was rotated similarly in the Look task, but in the Avoid task, it was rotated opposite to the cue. Since we observed inverted tuning in the Avoid task (higher responses away from the receptive field), the data was rotated to present the cue location opposite to the receptive field.

For tuning temporal cross-correlations, we adopted an approach previously used to remove auto-correlations from the data (Spaak et al., 2017). In this approach, data for the condition of interest (e.g., tuning in the Look task) was randomly divided into two halves, and the tuning function of one data half was correlated with the other data half.

We calculated mutual information for spatial cue locations during both the visual and memory periods. Mutual information refers to the amount of information that neurons carry about sensory and memory stimuli. A higher mutual information value indicates that the neuron’s activity carries more information about the stimulus location, implying a higher level of selectivity in encoding a specific location during sensory or memory periods (refer to Figure S10D). Mutual information for spatial locations was computed for each neuron and each task (Look and Avoid) separately. We equalized the number of trials per location for each neuron (for example, if the trial counts per location were [50, 55, 52, 53], we randomly selected 50 trials from each location without replacement). We repeated these mutual information calculations 50 times and then calculated the average of all iterations. During each iteration, we first computed the mutual information (“Observed mutual information”), and then we repeated the same procedure by shuffling the trial labels (“Shuffled mutual information”). Mutual information was defined as the difference between the observed and shuffled values.

#### Decoding of neuronal activity

We employed a linear Support Vector Machine (SVM) approach to discriminate between conditions based on high-dimensional neural activity profiles (Koren et al., 2020). All classification tasks were binary and conducted for each of the recording sessions (N=70), with the decoding accuracy subsequently averaged across these sessions. The input data for the classifier comprised neuronal spike counts averaged over a 200 ms period. During classification, the classifier was trained on 80% of the trials and then tested on the remaining 20%. Spike counts for both the training and test sets were normalized using Z-Score by employing the mean and standard deviation calculated from the training set. To ensure the robustness of our approach, we implemented a Monte-Carlo cross-validation strategy, involving 100 iterations with random divisions between the training and testing sets. The reported balanced accuracy was an average derived from these cross-validations.

The regularization parameter (C parameter) of the SVM was determined via a 10-fold cross-validation process performed on the training data. The parameter was selected from the following range: C > {0.001, 0.005, 0.01, 0.02, 0.05, 0.1, 0.5, 1, 10}. To assess the individual contributions of neurons in the classification process, we computed the decoding weights for each dimension (neuron) in relation to the classifier.

For the linear SVM constructed to decode the “cue in” vs. “cue out” conditions in the Look and Avoid tasks, we additionally permuted the trial labels for each iteration of the Monte-Carlo split. The mean balanced accuracy obtained from the label-shuffled dataset was used as a reference chance level to assess the statistical significance of our results.

## Acknowledgments

We are grateful to prof. Tirin Moore with mentorship and support for this project. Matthew Panichelo, Ruobing Xia and Warren Woodrich Pettine with help in data collection. Danielle Abreu Lopez, Shellie Hyde and Stephen Cital with technical assistance. This work was supported by NIH EY014924 and EY026877.

---

**Figure S1.**
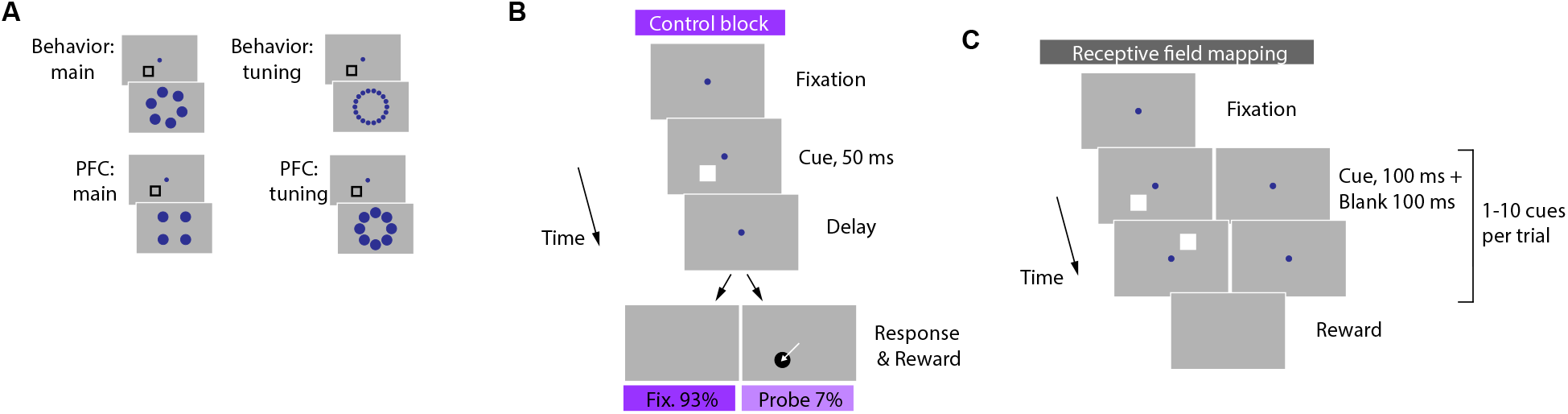
Methods. (**A**) Relative distances between two response targets in the experiments reported in this manuscript. Cue locations are rotated as if the cue always appeared at the shown location. Behavior: main - standard Look-Avoid tasks. Behavior: tuning - more distances between two targets were sampled (20 degrees apart). PFC: main - standard recording sessions in PFC. PFC: tuning - a small subset of recording sessions sampled more distances between two targets (45 degrees apart). (**B**) Fixation control task. A peripheral visual cue was task-irrelevant, and monkeys were rewarded for maintaining their gaze at the central fixation during the trial. Cue was the same as during the receptive field mapping task (next panel). On probe trials, after the delay, a single response target appeared, and monkeys made an eye movement to the probe location. (**C**) Receptive field mapping task. A peripheral cue was task-irrelevant, and monkeys were rewarded for maintaining their gaze at the central fixation during the trial. One to ten cues were presented during the trial.

**Figure S2.**
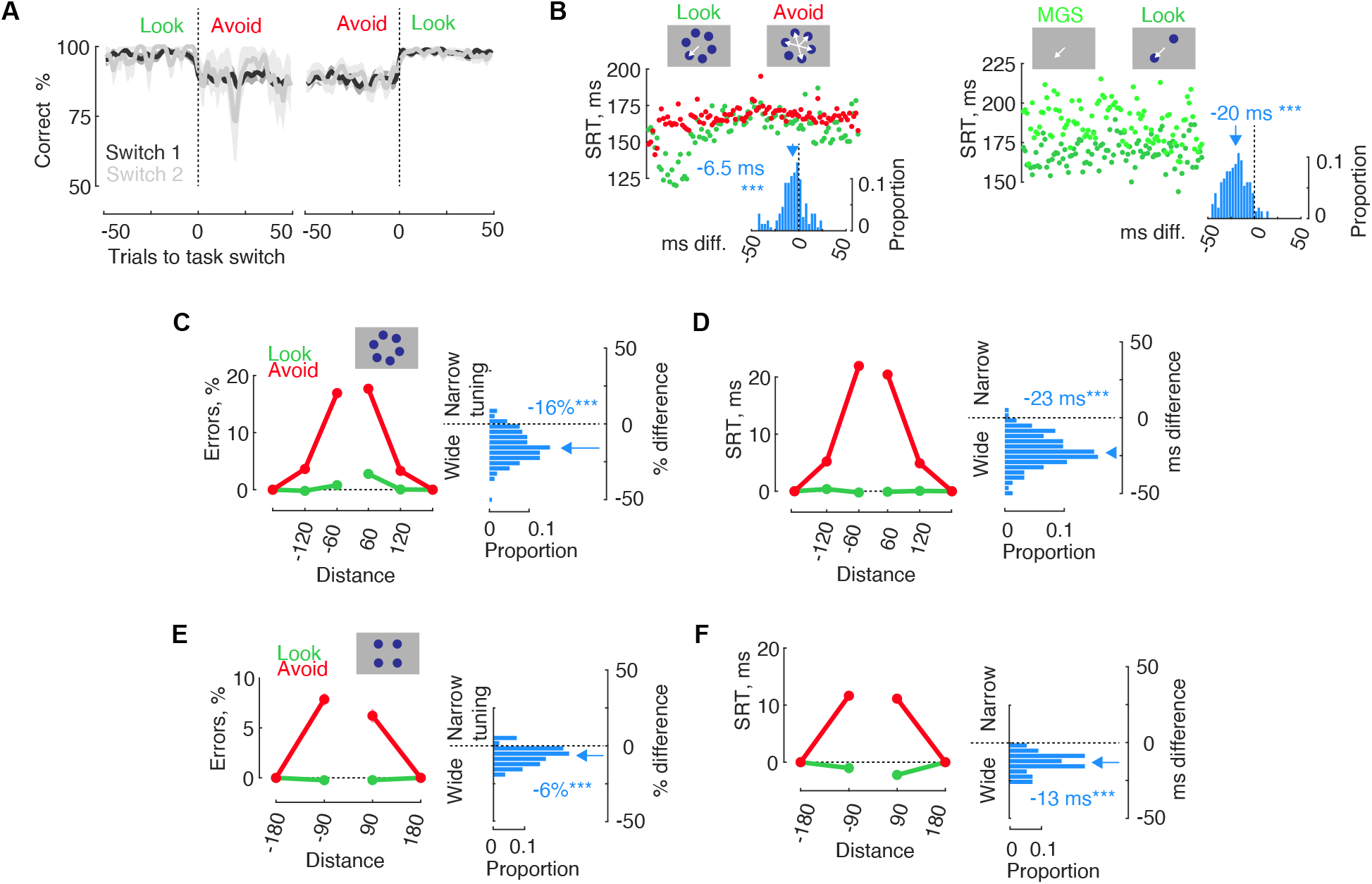
Monkey AQ performance. **(A)** Accurate memory task performance across multiple task switches. The task switched from Look to Avoid or from Avoid to Look at “Trial 0”. “1st switch” indicates the first switch from Avoid to Look on a given session. “2nd switch” represents the second switch on the same session (if completed). Performance remained stable before the switch, changed during the switch, and settled into a new stable state. We collected only a few 2nd switches from Look to Avoid (n=18) since monkey AQ typically did not perform the Avoid task late during the experimental session. (**B**) Reaction times in Look and Avoid tasks (left). Inset histogram - difference in reaction times between Look and Avoid tasks. Reaction times in Look and MGS trials (right) and the difference between the two. **(C)** Memory tuning in the main behavioral experiment (6 cue locations). Memory error probability is shown as a function of response target distances. Histogram insets: % memory errors difference between Look and Avoid, for individual behavioral sessions. Negative values indicate more errors or a wider tuning function in the Avoid task. (**D**) Tuning function constructed from saccadic reaction times on correct trials. (**E**) Memory tuning in the physiology experiment (4 cue locations). (**F**) Tuning function constructed from saccadic reaction times on correct trials.

**Figure S3.**
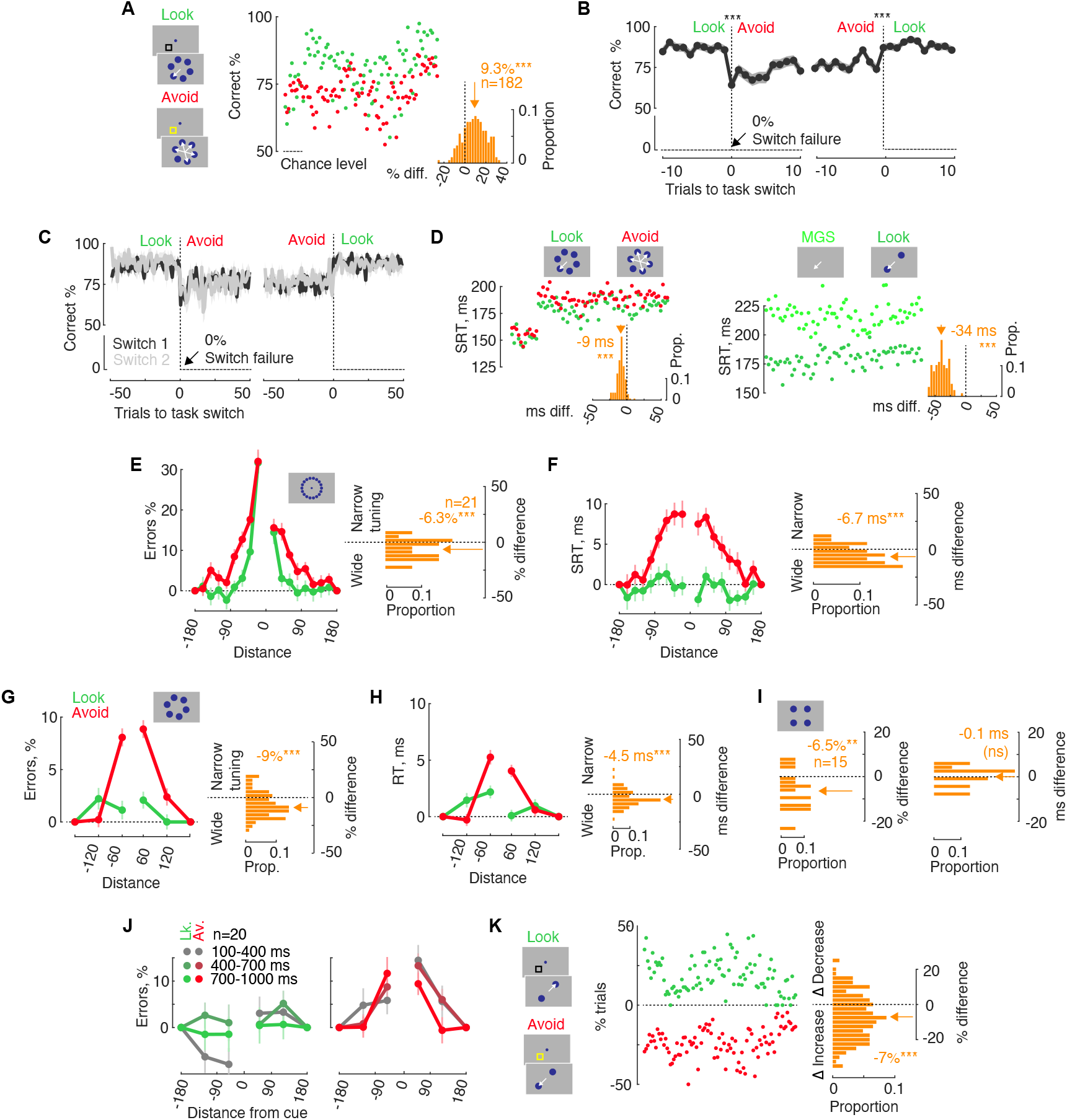
Performance, monkey HB. **(A)** Memory performance in Look and Avoid tasks. A randomly selected subset of 100 sessions, with each dot representing one session. Chance level is 50%. The inset displays the distribution of Look-Avoid performance differences for individual sessions. (**B**) Memory performance across task switches. The task switches from Look to Avoid or from Avoid to Look at “Trial 0”. We show 10 trials before and after the switch. (**C**) Memory performance across multiple task switches. The task switches from Look to Avoid or from Avoid to Look at “Trial 0”. “1st switch” refers to the first switch from Avoid to Look on a given session. “2nd switch” represents the second switch on the same session, if completed. (**D**) Reaction times in Look and Avoid tasks (left). Inset histogram - the difference in reaction times between Look and Avoid tasks. Reaction times in Look and MGS trials (right) and the difference between them. **(E)** Experiment with an extended range of response target distances (inset shows 18 possible cue locations). Memory error probability is shown as a function of response target distances. Histogram insets: % memory errors difference between Look and Avoid, for individual behavioral sessions. Negative values indicate more errors or a wider tuning function in the Avoid task. (**F**) Tuning function constructed from saccadic reaction times on correct trials. (**G-H**) Similar to (E-F), but for the main behavioral experiment with 6 possible cue locations. (**I**) Memory tuning differences during FEF recording sessions. (**J**) Experiment with an extended range of delay durations, randomly selected from an interval of 100 to 1000 ms. **(K)** Measure of failure to implement task rules. Performance errors are shown for trials when response targets were opposite from each other. Distribution of error probability differences between Look and Avoid tasks.

**Figure S4.**
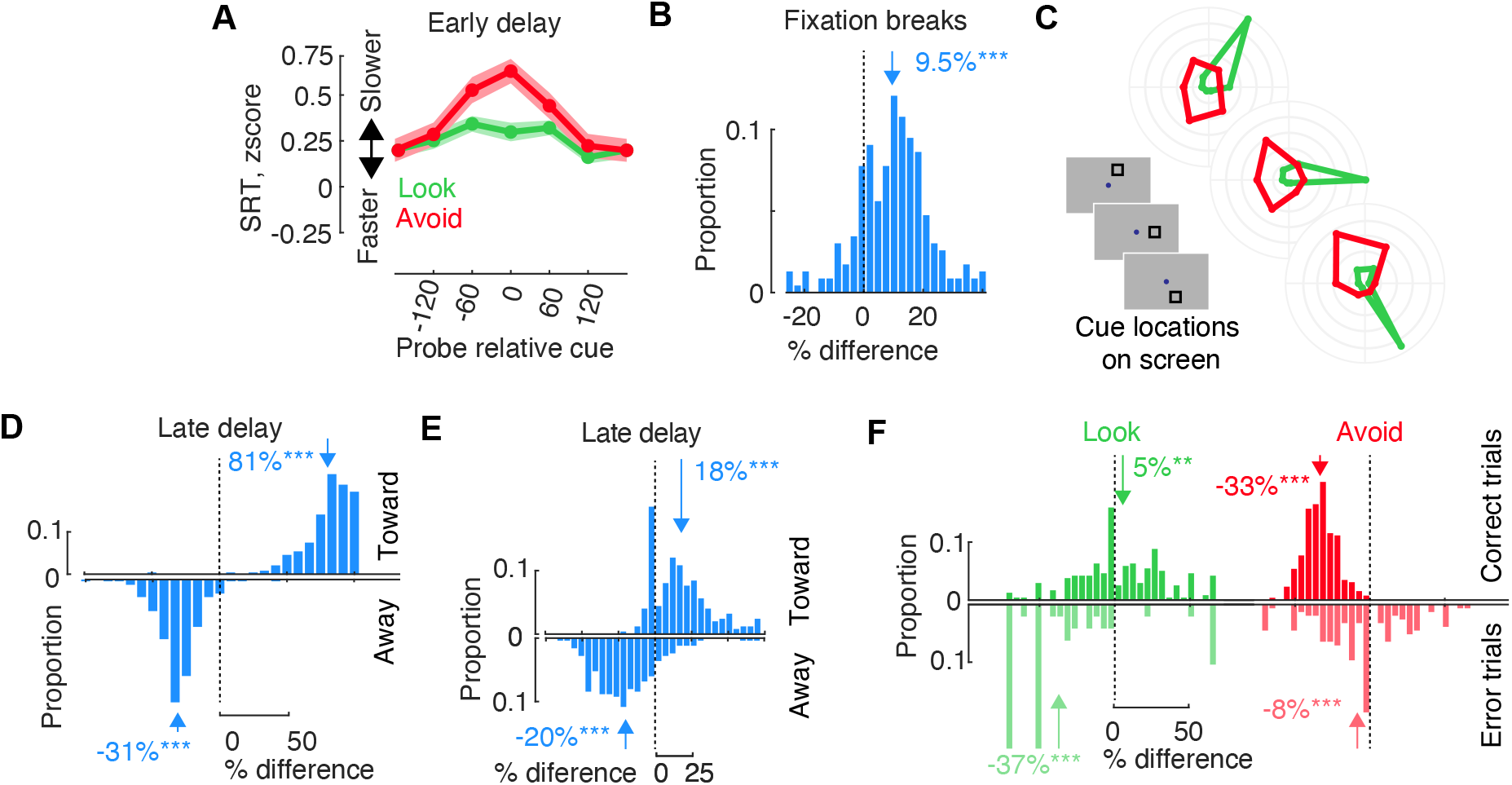
Sensory space and response space, monkey AQ. **(A)** Saccade reaction times during the early memory period (0-500 ms) as a function of probe location relative to the cue. Data is categorized for probes appearing in the first half and in the second half of the memory period. (**B**) Probability difference of fixation breaks during Look versus Avoid tasks. **(C)** Fixation break directions as a function of cue location on the screen. Insets depict the cue locations used in the task (up, right, down). The left and right cue locations are mirrored and combined. Polar plots illustrate fixation break directions on a given cue location (up, right, down). Fixation break directions vary in response to the cue location, indicating that they reveal memory location and the anticipated response. (**D**) Fixation break direction difference for Look versus Avoid tasks during the memory period. Histograms represent the Look-Avoid difference in “fixation breaks towards cue” and “away from cue” probability. Positive values indicate a higher probability of fixation breaks in the Look task. (**E**) Histogram of micro-saccade direction probability difference between Look and Avoid tasks during the late memory period. Histograms show the Look-Avoid difference in “micro-saccades towards cue” and “away from cue” probability. (**F**) The difference in “towards-away” micro-saccade probability during correct and error trials. Positive values indicate more micro-saccades towards the cue. Upper histograms represent correct trials, while lower histograms represent error trials.

**Figure S5.**
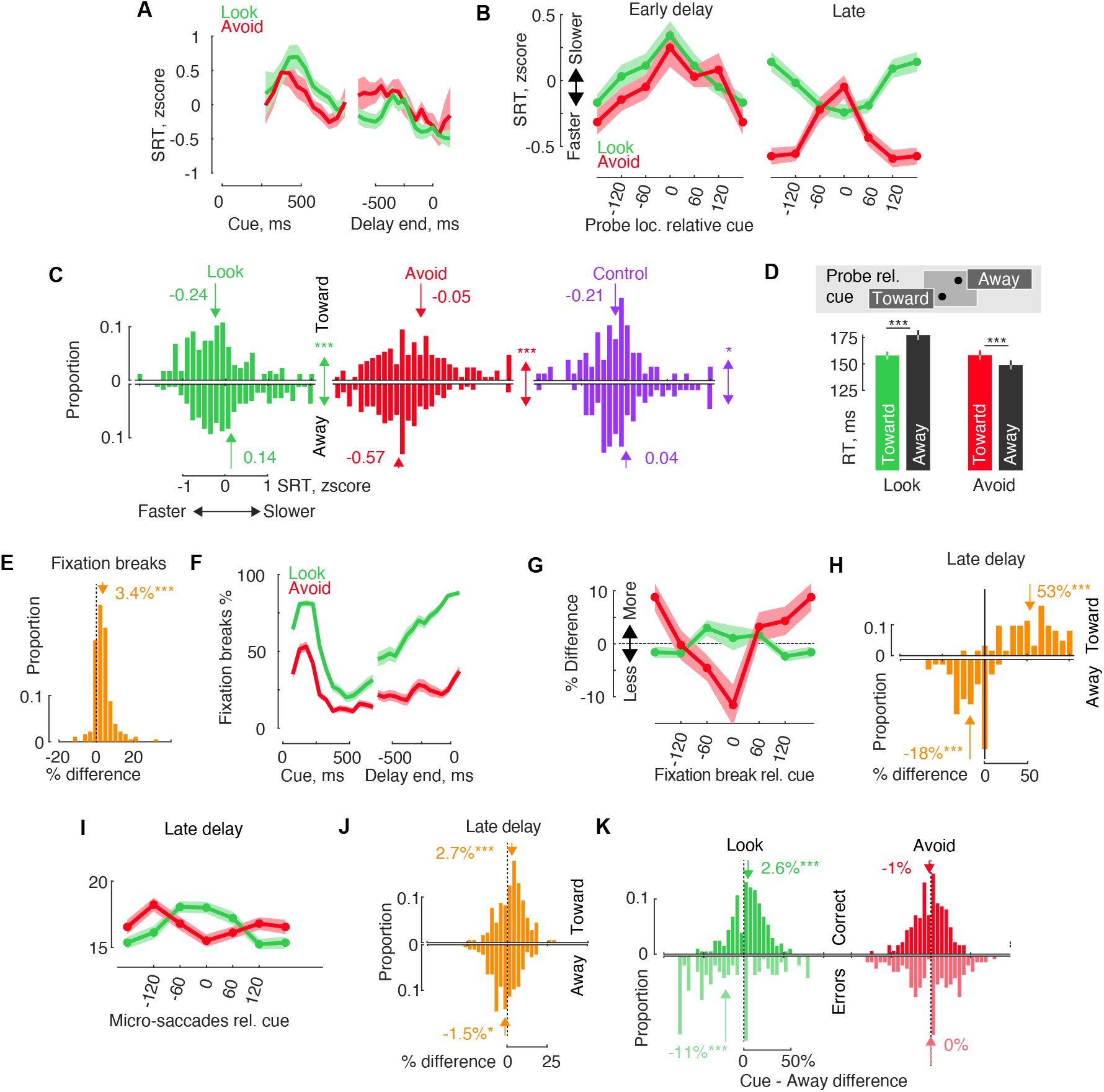
Sensory and action space, monkey HB. **(A)** Temporal course of saccadic reaction times to the cued location. Time is plotted as the stimulus-response onset asynchrony. **(B)** Saccade reaction times as a function of probe location relative to the cue. Data is categorized for probes appearing in the first half and in the second half of the memory period. The fixation task shows a small but significant facilitation pattern of results similar to the Look task, indicating a possible automatic action preparation towards the cue. (**C**) Histograms of individual trial reaction times for “probe toward” and “probe away” conditions in probe trials. A double-sided arrow indicates statistical comparisons between “probe toward” and “probe away” for each task. (**D**) Probe reaction times in neurophysiology recording sessions when only two probe locations relative to the cue were used. The inset illustrates probe locations relative to the cue. (**E**) Probability difference of fixation breaks during Look versus Avoid tasks. (**F**) Temporal course of fixation break probability toward the cue location. (**G**) Late-Early memory period difference in fixation break probability. Fixation break directions are relative to the cue location. Positive values indicate an increase in fixation breaks towards a location during the late delay, while negative values indicate a decrease in fixation breaks towards that location. (**H**) Fixation break direction difference for Look versus Avoid tasks during the late delay. Histograms represent the Look-Avoid difference in “fixation breaks towards cue” and “away from cue” probability. Positive values indicate a higher probability of fixation breaks in the Look task. (**I**) Micro-saccade direction probability relative to the cue location during the late delay. (**I**) Histogram of micro-saccade direction probability difference between Look and Avoid during the late memory period. Histograms represent the “micro-saccades towards cue” and “away from cue” probability. (**K**) Micro-saccade direction bias during correct and error trials. Bias is defined as the difference between the cue direction and the opposite direction. Positive values indicate more micro-saccades towards the cue. Upper histograms show correct trials, while lower histograms show error trials.

**Figure S6.**
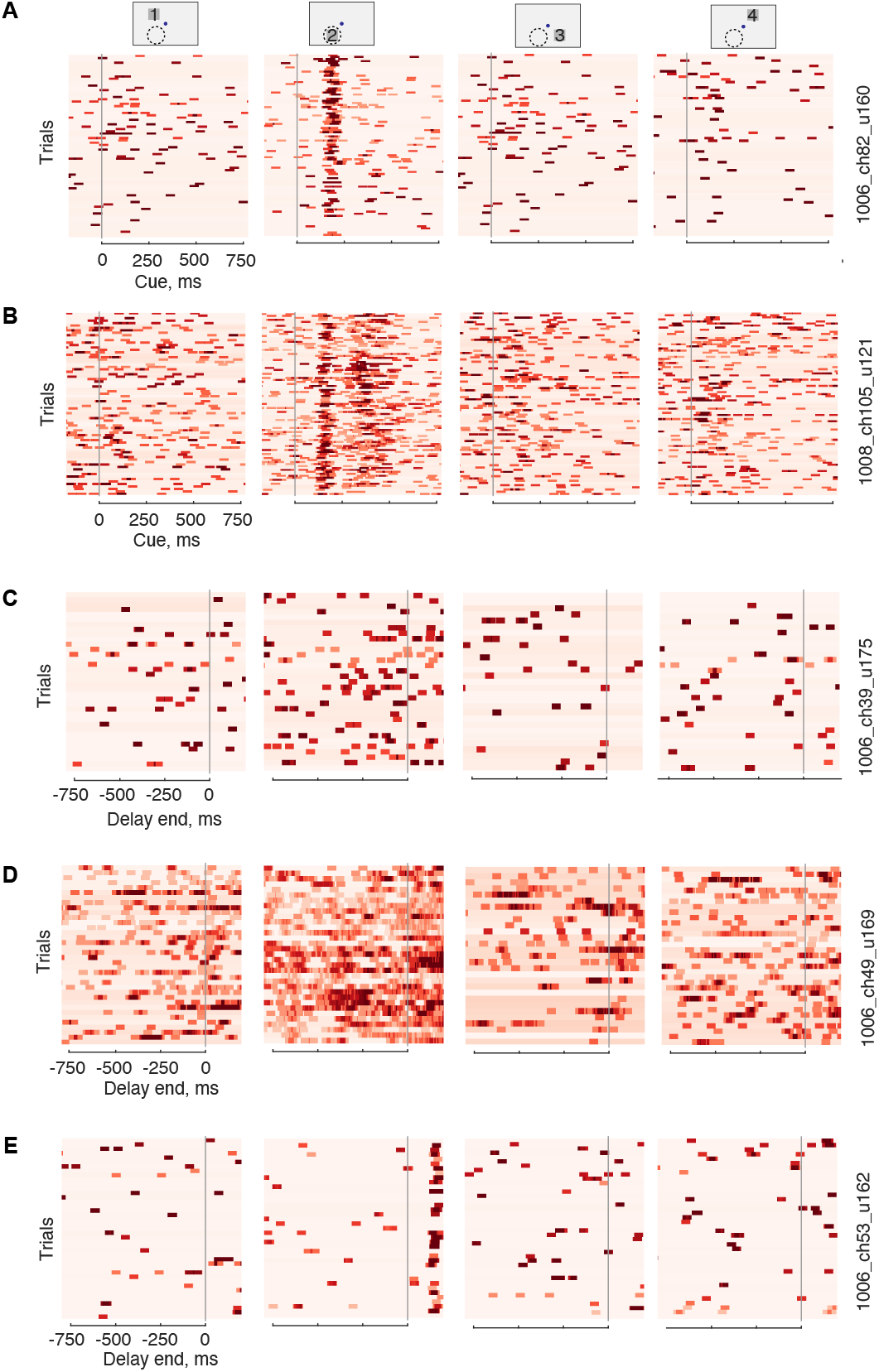
Example neuronal activity during different task periods. The activity for visual, memory, and target periods was determined independently for each unit. The inset indicates the cue location with a corresponding number, and the typical receptive field location is depicted as a dashed circle. **(A-B)** Detection of visual activity defined by ANOVA main effect of cue location 50-200 ms after the cue onset. The receptive field was detected at "Location 2". Firing rates are displayed using a 50 ms sliding window. These are Look trials, and each row represents one trial. **(C-D)** Detection of memory activity, defined by ANOVA main effect of memory location −300 to 0 ms before the end of the memory period. These are MGS trials. **(E)** An example unit without significant memory period activity, but with significant visual activity occurring after the memory. These are Avoid trials.

**Figure S7.**
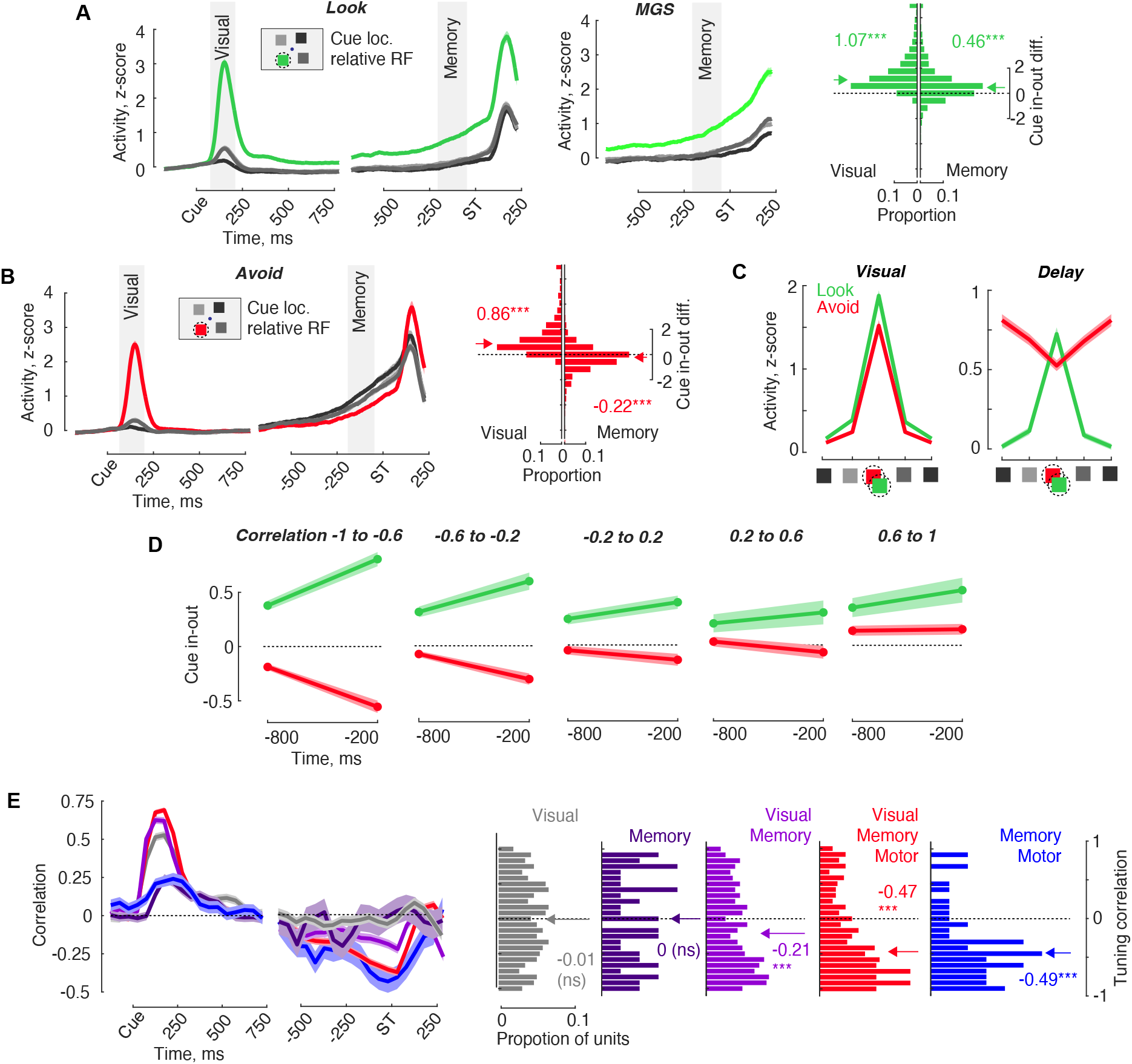
(**A**) Average population responses observed during Look and MGS tasks. Only units with significant memory activity are included. The activity is shown relative to cue onset and saccade target. The legend indicates cue locations relative to the receptive field (dashed circle). Shaded areas highlight the visual period (50 to 200 ms) and the delay period (−200 to −50 ms), which were used for data analysis. An inset illustrates the cue locations relative to the receptive field. The histogram displays the distribution of the difference between cues in the receptive field and cues opposite the receptive field for all neurons. (**B**) Population responses during the Avoid task. (**C**) Activity during the visual and memory periods of both Look and Avoid tasks. The activity is relative to the center of the visual receptive field. (**E**) Spatial selectivity - cue in-out difference during memory period −800 and −200 ms before memory delay end. Spatial selectivity is shown for groups determined by tuning correlation between Look and Avoid tasks during delay period (−200 to −50 ms) (Same as Figure 3). **(E)** The time course of correlations is depicted for units defined by significant visual, memory, and motor activity. The histograms show correlations during the delay period for units defined by visual, memory, and motor activity. The legend text for each histogram labels only periods of significant activity (“Visual,” “Memory,” “Motor”).

**Figure S8.**
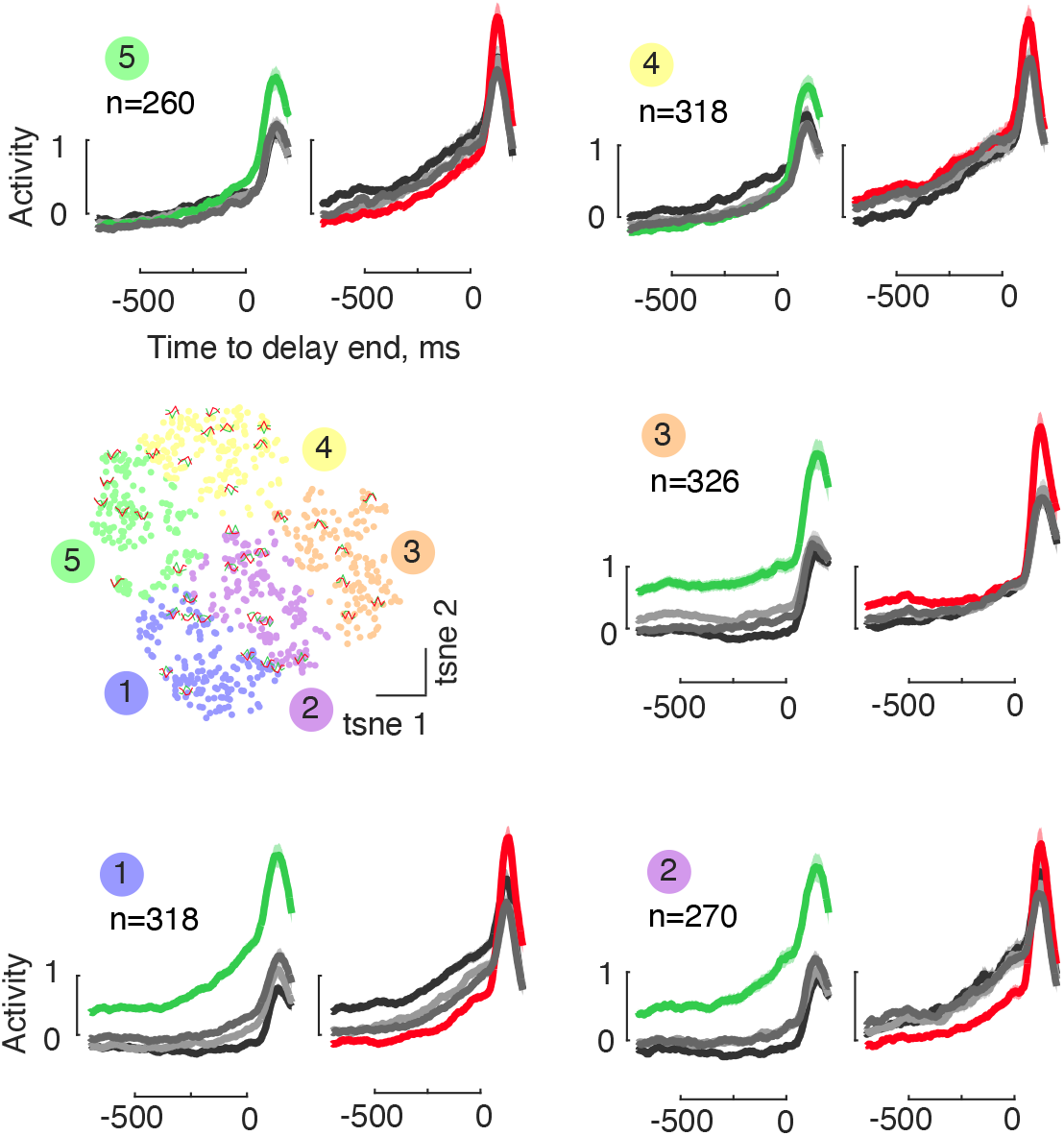
Clustering of Look-Avoid responses based on delay activity, using t-SNE space. Each neuron is represented by a color-coded dot in the scatter plot. Several randomly selected example tuning curves are overlaid on the dots. The average sub-population responses for each of the clusters in Look and Avoid tasks are displayed, with each cluster being identified by a number. In all clusters except one, the Look-Avoid task activity exhibited opposite tuning.

**Figure S9.**
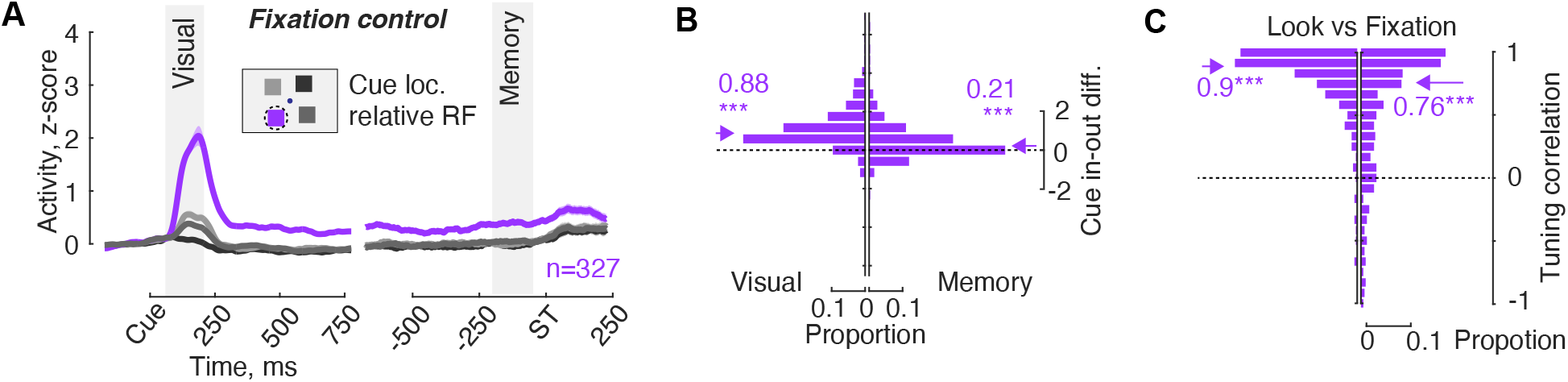
Fixation control task. (**A**) Average population responses in the Fixation control task, which was completed in a subset of sessions. In this task, monkeys maintained fixation during the trial. (**B**) A histogram displays the activity difference between the cue in the receptive field (RF) versus the opposite location. Positive values indicate higher activity for the cue in the RF. (**C**) Histograms illustrate the correlation between Look and Fixation tasks during the visual (left) and delay (right) periods.

**Figure S10.**
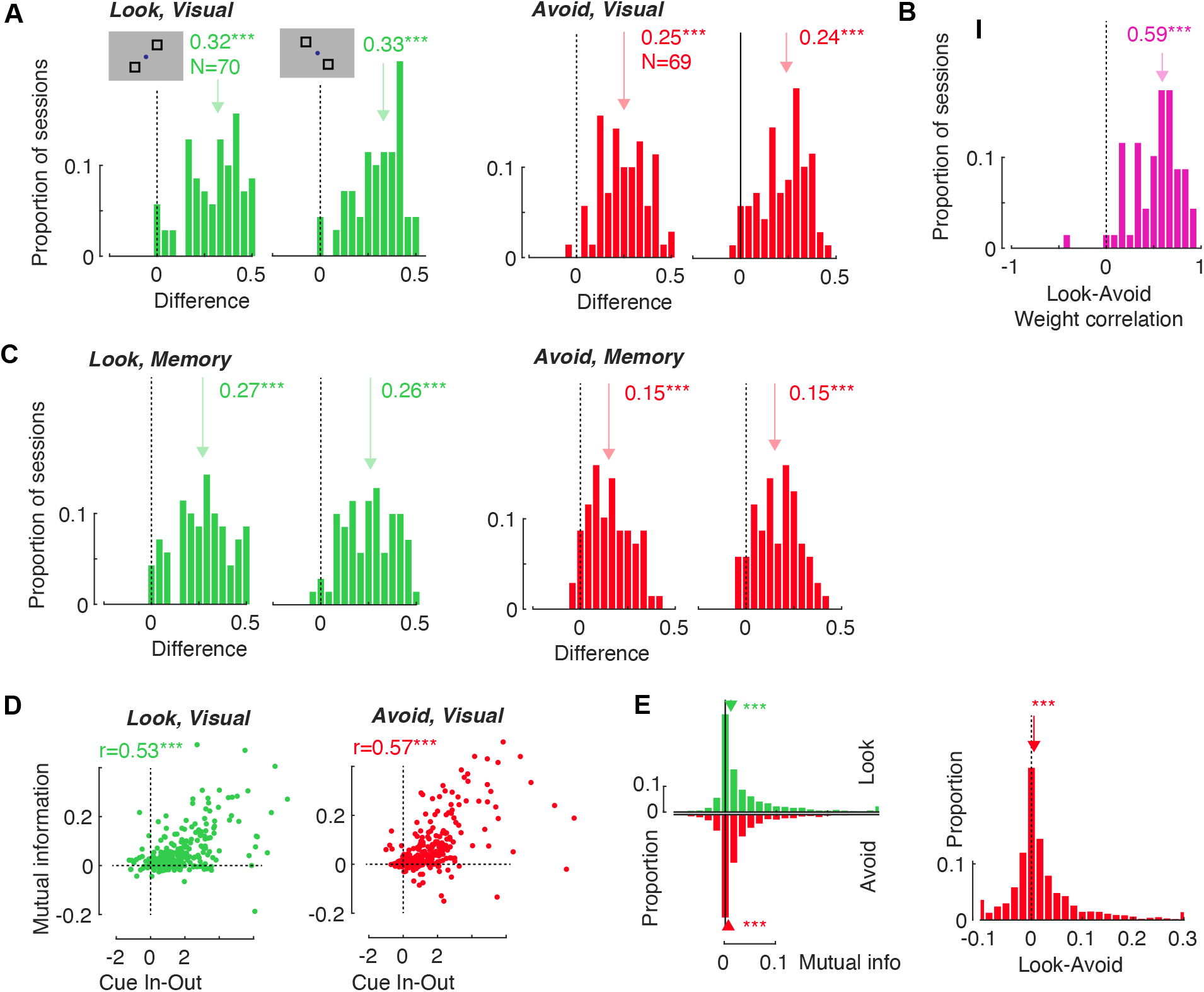
Bias and tuning during memory period. (**A**) The histogram depicts the decoding accuracy during the visual period of the task for both the Look and Avoid conditions. The decoder was trained using cue location combinations as displayed in the inset. For the final analysis, two sets of locations were merged. (**B**) Correlation of decoder weights between the Look and Avoid tasks during the visual period of the task. (**C**) Decoding accuracy during the memory period of the task. (D) The correlation between mutual information and neuronal activity associated with “cue in” versus “cue out” conditions. (**E**) On the left, the mutual information during the memory period. On the right, the difference in mutual information between the Look and Avoid tasks.

**Figure S11.**
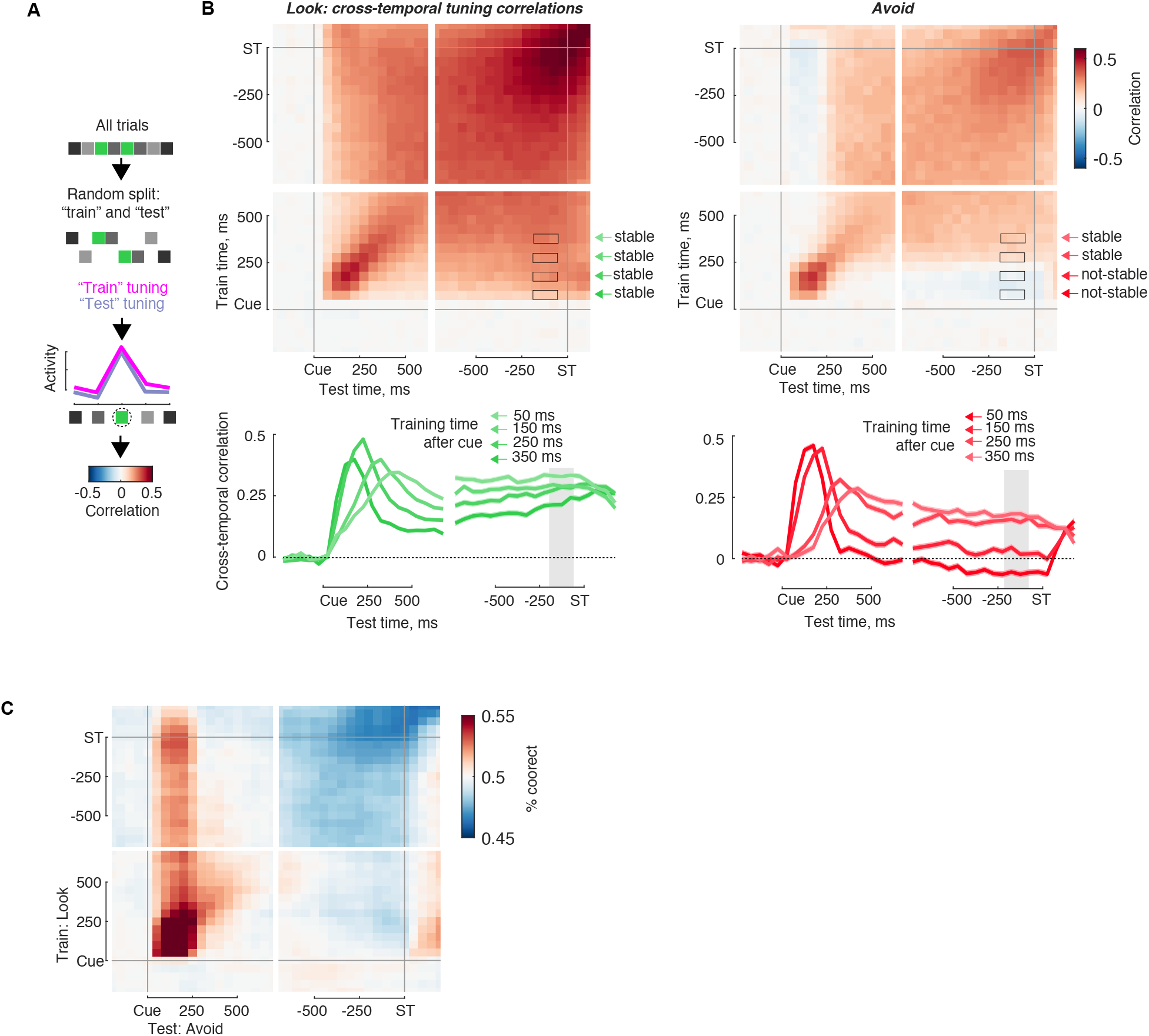
(**A**) Illustration of cross-temporal tuning correlation calculations. For each neuron, all trials were randomly assigned to “training” and “test” datasets, and cross-temporal tuning correlations were calculated between the training and test datasets. Positive correlations then indicate unchanged tuning across time. This approach removes auto-correlations along the diagonal. (**B**) Cross-temporal tuning correlations for Look and Avoid tasks. **(C)** Cross-temporal decoding between Look and Avoid tasks. Dataset was trained on Look activity, and tested on Avoid.

## Notes

### Competing Interest Statement

The authors have declared no competing interest.

